# How Changes in Common Practice Can Improve the Quality of Biomedical Science

**DOI:** 10.1101/218867

**Authors:** Matthias Steinfath, Silvia Vogl, Norman Violet, Franziska Schwarz, Hans Mielke, Thomas Selhorst, Matthias Greiner, Gilbert Schöenfelder

## Abstract

Assessing the impact of recommendations rectifying the reproducibility crisis (publishing both positive and “negative” results and increasing statistical power) on competing objectives, such as discovering causal relationships, avoiding publishing false positive results, and reducing resource consumption, we developed a new probabilistic model. Our model quantifies the impact of each single suggestion for an individual study and their relation and consequences for the entire research process. We can prove that higher-powered experiments can save resources in the overall research process without generating excess false positives. The better the quality of the pre-study information and its exploitation, the more likely this beneficial effect is to occur. Additionally, we quantify the adverse effects of neglecting good practices in the design and conduct of hypotheses-based research, and the responsibility to publish “negative” findings. Our contribution is a plea for adherence to or reinforcement of the good scientific practice and publication of “negative” findings.

## Introduction

A recent study (Freedman, Cockburn, and Simcoe 2015) estimated that the irreproducibility of published scientific data ranges from 51% to 89%. This is underlined by the outcome of a survey conducted by the *Nature* journal (Baker 2016). Here, the scientific community assumes that the reproducibility of published scientific studies is less than expected and desired (Baker 2016). Specifically, 90% of 1,576 scientist respondents agreed there is a significant or slight reproducibility crisis (Baker 2016). Interestingly, although published data cannot be reproduced, 31% of the respondents still do not believe that the published results could be wrong.

However, a lack of reproducibility often impairs the credibility of science and causes severe concerns of resource waste (e.g., money, time, human and animal resources, etc.), hindrance of the scientific progress and of new medical therapies, as well as the unnecessary suffering of experimental animals and wasting of a large number of animal lives, since many of the translational and preclinical studies in biomedical research are based on animal experiments. Therefore, the scientific community aims to find a solution for the reproducibility crisis.

There are many reasons why a study cannot be reproduced. Of course, an insufficient methodology description subsequently leads to irreproducible experiments and data, which is often justified by having the methods section relegated to small paragraphs in broad spectrum journals (Marcus and the whole Cell team 2016). Consequently, the methods sections often do not contain crucial information for reviewers or readers to evaluate the strength of the experimental data and draw conclusions (Marcus and the whole Cell team 2016). As such, simply reporting on experimental design and methods has to be improved to avoid reproducibility and robustness imbalances in science overall, with several publishers starting to reconsider how to report experiments. For example, Cell Press campaigned to empowering the methods sections (Marcus and the whole Cell team 2016).

More importantly, when an alleged effect (e.g., cause-effect relationship) either does not have the indicated strength or does not exist and, consequently, data cannot be reproduced although the experiment can be repeated. In this context, reproducibility is defined as the ability to exactly repeat an experiment and obtaining identical result within statistical error tolerances. Hence, the reverse conclusion is leading to the very relevant question: why a result cannot be reproduced if a published experiment can be repeated exactly?

Ioannidis identified a number of reasons that contribute to the non-reproducibility of most published studies in 2005 (Ioannidis 2005). These reasons are currently intensively discussed by the scientific community, and several practical recommendations were derived to find the way out of the “reproducibility crisis” (Munafò et al. 2017). We summarize these recommendations as follows:

A. positive results from studies indicate an effect, but “negative”/null results show no evidence of the expected effect. The positive predictive value (PPV) is defined as the probability that a positive prediction (an effect) is true. This quantity is deduced from the pre-study probability, and alpha and beta errors. A poor selection of hypotheses (effects) to be tested leads to a low degree of pre-study probability and, thus, decreases the PPV, the number of false positive results increasing. Consequently, one should carefully select the hypotheses to be tested.
B. a major problem addressed in literature (Munafò et al. 2017) is flexible design (e.g., differences between randomized versus observational trials not properly considered when a study is planned) and flexible data analysis for experiments (e.g., alternative approaches or methods exist for analyzing the same data), which increase the tendency (bias) for positive results. It is assumed that the proportion of statistically significant results is larger than anticipated when experiments are rigorously conducted. This problem was theoretically investigated, but is also confirmed by empirical studies and meta-research (e.g., in neurosciences, the amount of statistically significant results exceeds any rational expectation) (Tsilidis et al. 2013). Systematic errors in the statistical evaluation of experiments in neuroscience are one reason that could lead to a bias in favor of the positive results (Kriegeskorte et al. 2009, Nieuwenhuis, Forstmann, and Wagenmakers 2011). Another reason is the consequence and prevalence of attrition in preclinical research through biased animal removal (Holman et al. 2016). In fact, the distribution of published p-values hints at a bias due to “P –hacking,” which means the data are produced or analyzed until the p-values fall below the significance level (Head et al. 2015). Results obtained from such flawed experiments are therefore biased. Prospective, rigorous, and transparent experimental design, and blinding and randomization of studies are proposed prevention measures (Begley and Ioannidis 2015). The pre-registration of studies would also be an improvement (De Angelis et al. 2004, Simes 1986, Chambers 2013).
C. despite their potential, the probability of publishing “negative”/null results is lower than that for positive results (Tsilidis et al. 2013). For instance, “negative”/null results repeatedly remain in lab books, on hard disks, and drawers, but if reported at all, they are often published in low impact journals or as by-products of positive outcomes and are, hence, difficult to observe. Accordingly, a large number of research findings remain unnoticed. The consideration of such results is much lower for further studies and, thus, may not influence scientific progress. In summary, a lack of information about “negative”/null results leads to publication bias, because positive effects are overestimated (e.g., in meta-studies) and false-positive results are not disproved. To avoid these effects it is suggested to publish all “negative”/null results (Ioannidis 2006).
D. the power of the statistical tests used to analyze experiments depends mainly on sample size, which is equal to the number of animals in a study for animal experiments. Lower power means that the false negative rate increases, which results in a lower probability to find the true effects. Additionally, the probability of a positive finding to be true (i.e., PPV) is also decreased, because the PPV decreases as power decreases. Therefore, a prior sample analysis is recommended to achieve the appropriate power. As such, fewer but larger studies are recommended, because research findings become more likely true (Cressey 2015).

Indeed, all four recommendations sound very plausible to improving reproducibility in science, but have to be proven. Here, the gold standard is the empirical analysis of the recommendations’ impact after their implementation within the scientific community. However, this approach is likely to take years. Thus, mathematical models are a useful alternative approach for the proof of concept.

Models using evolutionary or game-theoretic approaches are helpful to understanding how the dynamic of the scientific cognitive progress is influenced by human conflict and cooperation within a competitive situation. In other words, these models help understand why currently “bad science” (i.e., all four aforementioned recommendations are not appropriately consider) still exist. For instance, Smaldino and McElreath (Smaldino and McElreath 2016) demonstrated that poor methodological practices and high false discovery rates increase how an incentive structure rewards publication. Even replication does not stop this process. According to this approach, research groups may survive (i.e., “evolve”) within the scientific community on the basis of the sheer number of publishable results, which may constitute an incentive for applying poor science. Additionally, research groups maximize their fitness, but not the scientific progress when spending most of their effort seeking novel results and conduct small studies that have only 10%–40% statistical power (Higginson and Munafo 2016). This was recently demonstrated by Higginson and Munafó (Higginson and Munafo 2016).

Other models (McElreath and Smaldino 2015, Nissen et al. 2016, Bakker, van Dijk, and Wicherts 2012) serve as a formal framework for reasoning about the normative structure of science. They help determine standards for research practice (e.g., sample sizes) and are used to optimize certain target variables (e.g., the PPV). While Bakker (Bakker, van Dijk, and Wicherts 2012) models the causes and effects of questionable research that leads to biased effect strengths, two models with similar structure (McElreath and Smaldino 2015, Nissen et al. 2016) have investigated the impact of both the likelihood to replicate an experiment and of publishing “negative”/null results on the quality of science. Whereas Nissen et al. (Nissen et al. 2016) focused on a single claim at a time, McElreath and Smaldino (McElreath and Smaldino 2015) analyzed the progress of a group of scientists testing a suite of hypotheses. Both models were motivated by the concern that numerous published research findings are false, as pointed out by Ioannidis (Ioannidis 2005). Although both models already take into account the selective publication of positive results, we are persuaded that both models are not sufficient to provide enough information on the impact of the four aforementioned recommendations. We are convinced that two relevant aspects of the scientific process are not well captured in the above-mentioned approaches:

I. Published experimental results will cause or forestall further experiments concerning the same research problem (“bad science paves the way for more bad science”).
II. Because of limited resources (e.g., number of laboratory animals or study participants) the model should explicitly consider the consumption of various resources, most importantly the use of experimental animals (“bad science is a waste of laboratory animals”).

Considering these two aspects is especially important, as the impact of the recommendation should be measured on three objectives: A) uncovering the true causal connection (sensitivity; e.g., the etiology of a disease to be discovered), B) avoiding false positive results (specificity), and C) minimization of resources in a single experiment (e.g., number of animals in a single experiment) (Bate and Clark 2014, Erb 1990). It is also important to test how the individual recommendations effect all three objectives simultaneously.

Therefore, we proposed a novel model of the research and publication process, which exceeds the setting of a single experiment, to evaluate the impact of the aforementioned recommendations on the overall scientific process.

## Methods

### Model

The entire process of hypotheses testing research, encompassing consecutive studies and publications, is described in our probabilistic model. To better explain our model, we start with the simple case (initial model) of only one research team that explores an observed effect, such as the triggering of a signaling cascade by a certain small molecule. Due to published or own pre-study experiments, the research team has two probable scientific hypotheses: the small molecule stimulates receptor A or inhibits receptor B. Based on pre-study knowledge, one of the hypotheses is more likely true than the other. The research team will now begin to statistically plan an experiment that covers the scientific hypothesis more likely true (i.e. the team plans to investigate the factor (F_1_) that is most likely causal). In other words, the most likely hypothesis will be tested first. Let us assume this is the scientific hypothesis that the small molecule stimulates receptor A. To calculate the sample size needed for the experiment, the team decides for significance level α and statistical power (1 - β), given the expected effect size δ. After completing the experiment, the statistical test leads to either rejection (small molecule stimulates receptor A) or acceptance (small molecule does not stimulate receptor A) of the null hypothesis. For simplification, we will henceforth call the rejection of the null hypothesis “positive result” and the acceptance “negative result.”

As the outcome of a statistical test can either be true or false, there are four different possible results: true positive, false positive, true negative, and false negative (Figure 1). These four results lead to different overall outcomes for the research problem (i.e., answering the question of the mechanism by which the small molecule triggers the signaling cascade). In case of a true positive result, the research problem is solved and no further trials are necessary, that is, the team will not test their second hypothesis that the small molecule inhibits receptor B. The research team will, however, publish their results that the small molecule stimulates receptor A.

**Figure 1:**
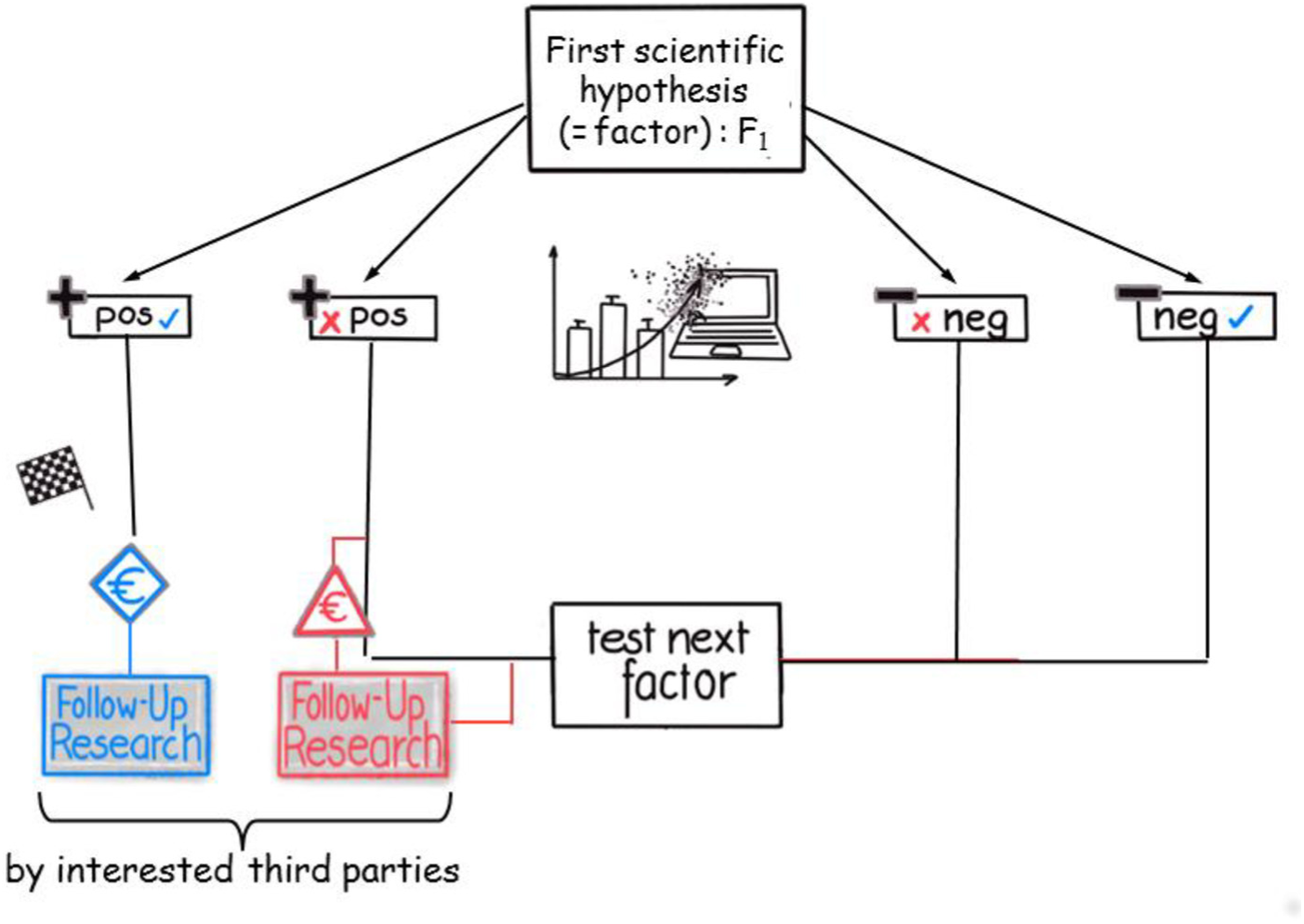
Flow diagram of the modeled research and publication process (1) Different possible sequences of outcomes, actions, and decisions are displayed. One research team investigates the first factor, F_1_. After experiment completion, the statistical test yields positive (left side) or “negative”/null results (right side). These results can either be true (blue checkmark) or false (red cross). Positive results are always published. In the case of true positive results, the research problem is solved and third parties can (successfully) use the results, while false positive results can be misleading for third parties, costly, and possibly dangerously. “Negative”/null results lead to an exclusion of the factor from further investigations; as a consequence, factor F_2_ is tested next. In case of a false negative result, this true (causal) factor is overlooked and, after the investigation of all remaining factors, the research process ends without a discovery.

For a false positive result, the research team will react similarly. The team believes that the research problem is solved and will publish the result. We assume that all positive results are published. Furthermore, we assume the research problem and its results are interesting or relevant enough to be noticed and used by third parties. For instance, there might be researchers looking for substances that stimulate receptor A, and they use the small molecule as positive control in an experiment.

In the long run, these assumptions lead to an important difference between a true positive and a false positive published result: the more additional research is based on or uses the true positive result, the more it will become apparent that the original hypothesis holds (verification of a true positive result, which is always the case in our model). The reverse is true for the false positive result, but the cost of identifying a positive result as false might be immense and—in case of animal tests or even clinical trials—not only financially, as humans or animals could be harmed in the process. The original research team will eventually realize that their originally published positive result was a false positive and will test the next hypothesis (i.e., the small molecule inhibits receptor B). In conclusion, we assume a false positive result is always falsified, which means that a false positive result is eventually detected, because a variety of different experiments (e.g., follow-up research by other teams) based on the published false positive result will fail. See supplement file SI2 for removal of these restrictions.

Our example research team will continue to test its second hypothesis if the original statistical test after their first experiment gave a true or false negative result: the detection of a true negative result will allow the research team to test the second—possibly true—hypothesis to find the solution to its research problem. As there is only this one research team working on this research question, publishing the negative result will have no impact. A false negative result will eventually lead to the end of the research process (all available hypotheses were tested) without discovery, because the true first hypothesis was rejected and the false second hypothesis will be also rejected (even a possible false positive result will ultimately be uncovered due to research based on the said result (falsified), as explained above). The entire course of action is depicted in Figure 3.

Most parameters we use to model the process of hypotheses testing research can be explained with this simple case of one research team with two hypotheses. First, based on pre-study information, the research team identifies a number of potential factors (in our example, two hypotheses: I) the small molecule stimulates receptor A or II) inhibits receptor B) and ranks both — the one that is likely true is tested first. However, this pre-study information can be comprehensive or scarce, and its availability, usage, or interpretation to create pre-study knowledge can be more or less biased. To model this, we need to include different qualities of pre-study information in a quantitative and objective manner. Consequently, we introduce pre-study probabilities ω_i_ for each hypothesis *i*, which are governed by the parameter π_k_. Figure 2 depicts an example with three factors (i.e., three hypotheses).

**Figure 2:**
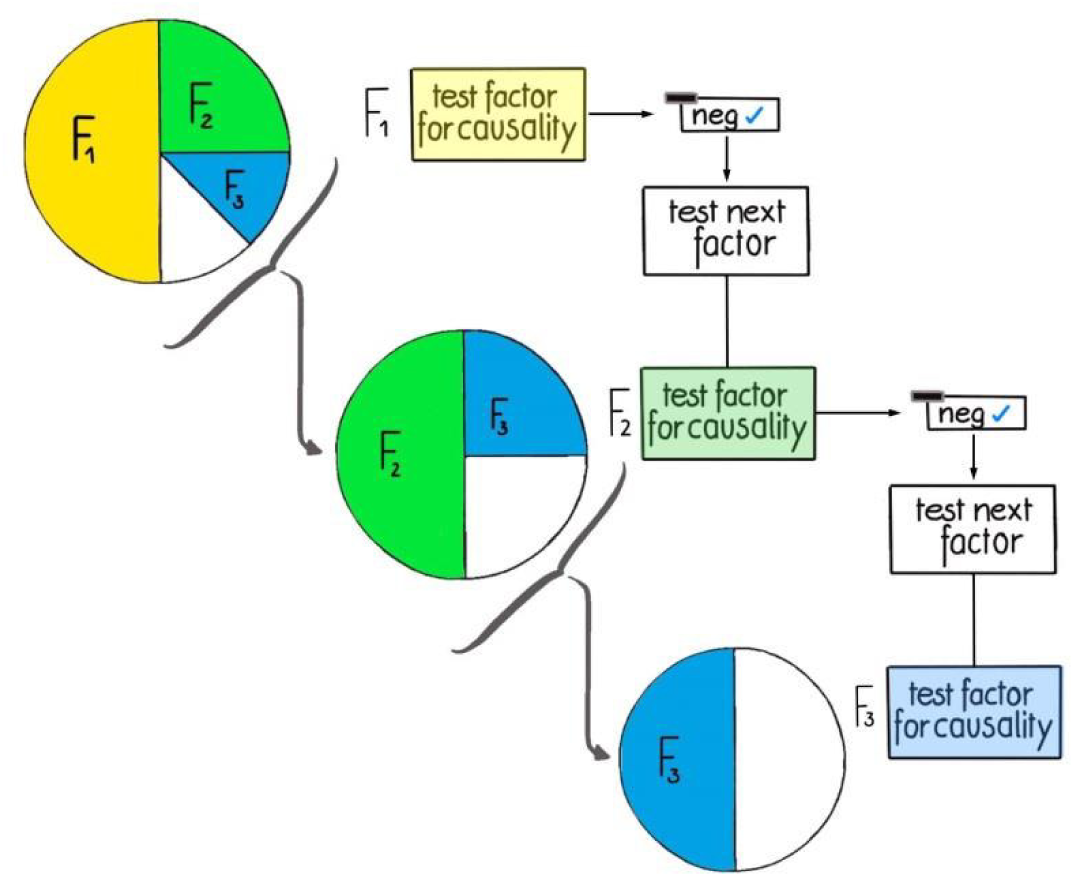
Figurative representation of how π_k_ defines pre-study probabilities. With π_k_= 0.5, the pre-study probabilities of F_1_, F_2_, or F_3_ to be causal are 50%, 25%, and 12.5%, respectively. If, however, F_1_ is tested and found to be not causal, the probabilities for the remaining factors will increase, the second factor F_2_ will become the most probable one, with a pre-study probability of 50%, and the pre-study probability for F_3_ will be 25%, etc.

The higher π_k_ the higher the probability that the first factor in the ranking is the “right” (i.e., causal) factor. In other words, according to the formula given in the supplement (SI1) (equation 1): if π_k_ = 0.5, the pre-study probability of the first factor (ω_1_) to be causal is 50%, if π_k_= 0.9, then ω_1_= 90%. To simplify the modeling procedure, π_k_ is defined as to maintain the pre-study probability ratio between two consecutive ranks constant. This is depicted in Figure 2 (for more details, see the supplementary information (SI1). Therefore, π_k_ can be interpreted as the quality (i.e., predictive power) of the pre-study information available for the research problem under investigation.

Most importantly, in reality, even if there is high quality pre-study information, circumstances such as current research trends, financial considerations, availability of publications, and also the “human factor,” lead to situations in which this existing information is not fully exploited, leading to incomplete pre-study knowledge. Basically, this can result in the incorrect prioritization of research activities. Our model covers this by including the possibility that factors might be tested in a “wrong” order (i.e., not in the order defined by their pre-study probabilities). Using our simple example, this means the research team first investigates the hypothesis that the small molecule inhibits receptor B (second hypothesis, F_2_, less likely to be true), because receptor B seems more interesting.

The research team statistically plans the first experiment, calculating the needed sample size by setting the significance level α, and the statistical power (1 - β). Additionally, the resulting sample size also depends on the effect size δ they estimate based on pre-study information. For simplicity, in our model, the same sample size calculation is used for all factors for one research problem, and all factors considered for a research problem have either zero effect strength (non-causal factors) or the same finite effect strength (causal factors).

Subsequently, there is one probabilistic parameter our model uses that is only needed for more than one research team investigating a research question. This is the probability to publish “negative”/null results P_pub_; as previously mentioned, such publications are irrelevant if there is only one research team involved. Let us assume a second team is also interested in the said research question. However, this second team may be not as fast as the first team in preparing the experiment, perhaps because for the second research team another problem has priority. If the first team publishes a positive result for receptor A, the second team will not repeat the same experiment. In contrast, if the first team tests receptor A negatively but does not publish the result, the second team will investigate receptor A, which means more resources are used. The same procedure is repeated with each of the subsequent hypotheses tests by the first group. The procedure for more than two teams is analogously defined. Therefore, the number of teams, n_t_, involved in solving a research problem is an important parameter for the total amount of used resources (and wasted time).

Additionally, we considered another scenario. In parallel to the investigation of receptor A by the first research team, a second research team decides to investigate receptor B–, even if it is aware that receptor B is less likely to be causal. Therefore, the number of factors tested in parallel by the different teams, n_p_, is an additional parameter (this scenario is described in more detail in the supplement (SI2)).

In our model, we vary β and P_pub_ to quantify their impacts on the overall number of true positives (which designates the scientific gain, g), the overall number of false positive results fp, and the amount of laboratory resources needed for the entire research process (in the following, we use the total number of samples needed (n_total_) as a measurement for those resources; supplement equations (SI1)15, 17, and 18). For clarity, since this study is about statistics and probabilities, the expectation values, that is, their theoretical mean values, are actually considered and denoted as E{g}, E{fp}, and E{n_total_}. For example, considering all probabilities, in scenarios with only one true hypothesis, the scientific gain E{g} is a real number between 0 and 1 and describes the probability to obtain the true positive result.

While the model basically postulates that the rules of good scientific practice (GSP) are followed, it also includes the possibility to deviate from GSP and thereby increase the possibility of positive results. To model these deviations we introduce parameter u, which is defined in the same way as by Ioannides (Ioannidis 2005): the *“proportion of probed analyses that would not have been “research findings,” but nevertheless end up presented and reported as such, because of bias.”* Furthermore, we describe and apply another approach in the supplement (SI2), which resembles the questionable research practices described by Bakker et al. (Bakker, van Dijk, and Wicherts 2012). As previously mentioned, we include another form of bias in our model: we postulated that the hypotheses are not tested in the order of declining pre-study probabilities, but in a completely different order (e.g., due to hypes or available technology).

**Figure 3:**
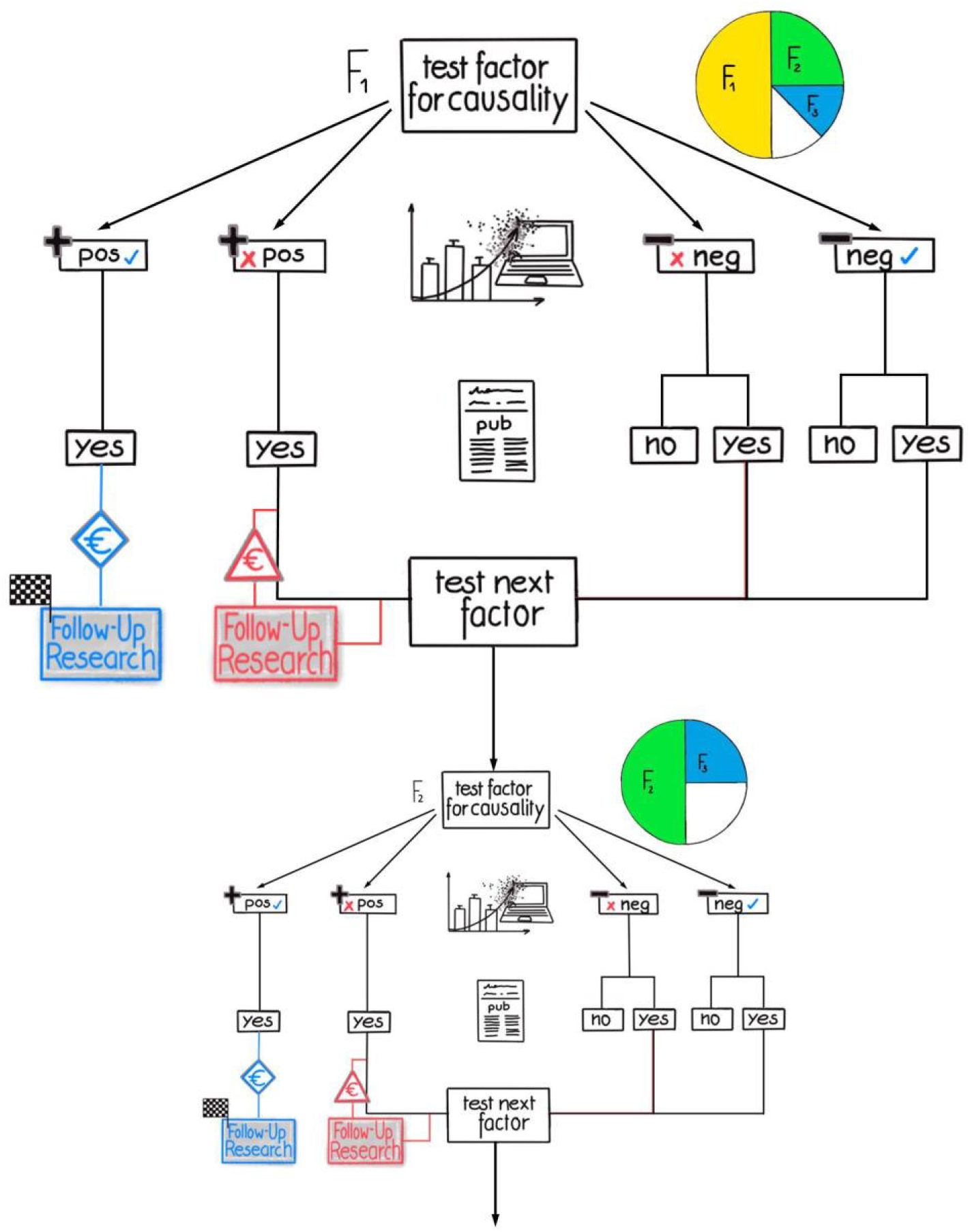
Flow diagram of the modeled research and publication process (2)

The different possible sequences of outcomes, actions, and decisions are displayed when several research teams are involved and “negative”/null results are published with probability P_pub_ (see Figure 1): published “negative”/null results have an influence on “competing” research teams so that those factors do not have to be tested more than once.

### Results

In this paper, we focus on the effects of statistical power (1 - β) and probability to publish “negative”/null results (P_pub_) on the scientific gain (g), number of false positives (fp), and total number of samples (n_total_) required throughout the entire research process. Additionally, the consequences of scientifically biased experimental approaches are investigated by introducing and varying parameter u (Ioannidis 2005). This parameter defines the proportion of experiments, which would have a negative test result in case of an unbiased procedure, but will present as positive because of a biased procedure (Ioannidis 2005). We also examined scenarios in which the potential factors are tested in a “wrong” or biased order. As stated in the previous section, all effects on g, fp, and n_total_ are determined by calculating their corresponding expected values (E{g}, E{fp}, and E{n_total_}).

#### Systematic analysis of the influence of statistical power

Here, we are especially interested in the question of whether the competition between our three objectives can be partially overcome. Firstly, let us examine significance level α. In the initial model, α has an impact only on both the number of false positives E{fp} and the selection of sample size n_a_, but in a different direction: while E{fp} increases with α, the sample size n_a_ decreases with α. Consequently, either E{fp} or E{n_total_}) can be minimized when only α is varied. The solution is to choose a value of α to limit the number of fp, hold it constant, and then vary statistical power (1 - β) only to address this question properly. If not mentioned otherwise, a value of 0.05 is used.

The influence of statistical power (1 - β) on the reliability of experimental results in a single experiment is well known. The higher the power, the lower the possibility to miss a causal factor. We can show that this aspect holds true even if u and P_pub_ are considered. Hence, the statement applies for entire research processes: with an increasing power (1 - β) the scientific gain (E{g}) increases monotonically. This relationship can be proven with the derivative of the corresponding model equation (see supplement SI1).

It is also established that the false discovery rate, that is, the number of false positive results divided by the number of all positive results, decreases with (1 - β) (Button et al. 2013). Additionally, using the equations deduced from our model (see supplement SI1), we can show that even the absolute number of published false positive results (E{fp}) decreases during the research process. This means that, with an increase in power (1 - β), less false positive results will be obtained. Therefore, in this framework, there is no conflict between optimizing E{g} and E{fp}, and increasing the statistical power has only positive effects.

Once again, let us look at a single experiment where scientists calculate the matching sample size for a targeted statistical power. Here, an increase of statistical power always leads to a higher sample size. What happens if we regard the entire research process? Applied to our model, the answer is remarkable: E{n_total_} is not necessarily a monotonic function of β. The consequence of this statement is that, even if the sample size of a certain single experiment increases with the statistical power, the expected number of studies, and, therefore, the total number of samples E{n_total_} needed to solve the research problem can decrease. Hence, an intriguing question arises: are there useful local or global minima of E{n_total_(β)} so that there is a value β_min_ that minimizes the total number of samples needed to find the true causal factor, without generating an excess of false positives? To answer this question, we have to examine the possible scenarios for E{n_total_(β)}. Some conclusive examples can be found in Figure 4. The left-hand diagram (panel A) compares different combinations of values for P_pub_ and bias u. The right-hand diagram (panel B) illustrates the consequences of that type of bias, where the order in which the factors under investigation are examined is changed. In both panels, the lower line (dark blue) represents the scenario, in which all results are unbiased and published (P_pub_= 1, u = 0) and are obtained in the correct order.

**Figure 4:**
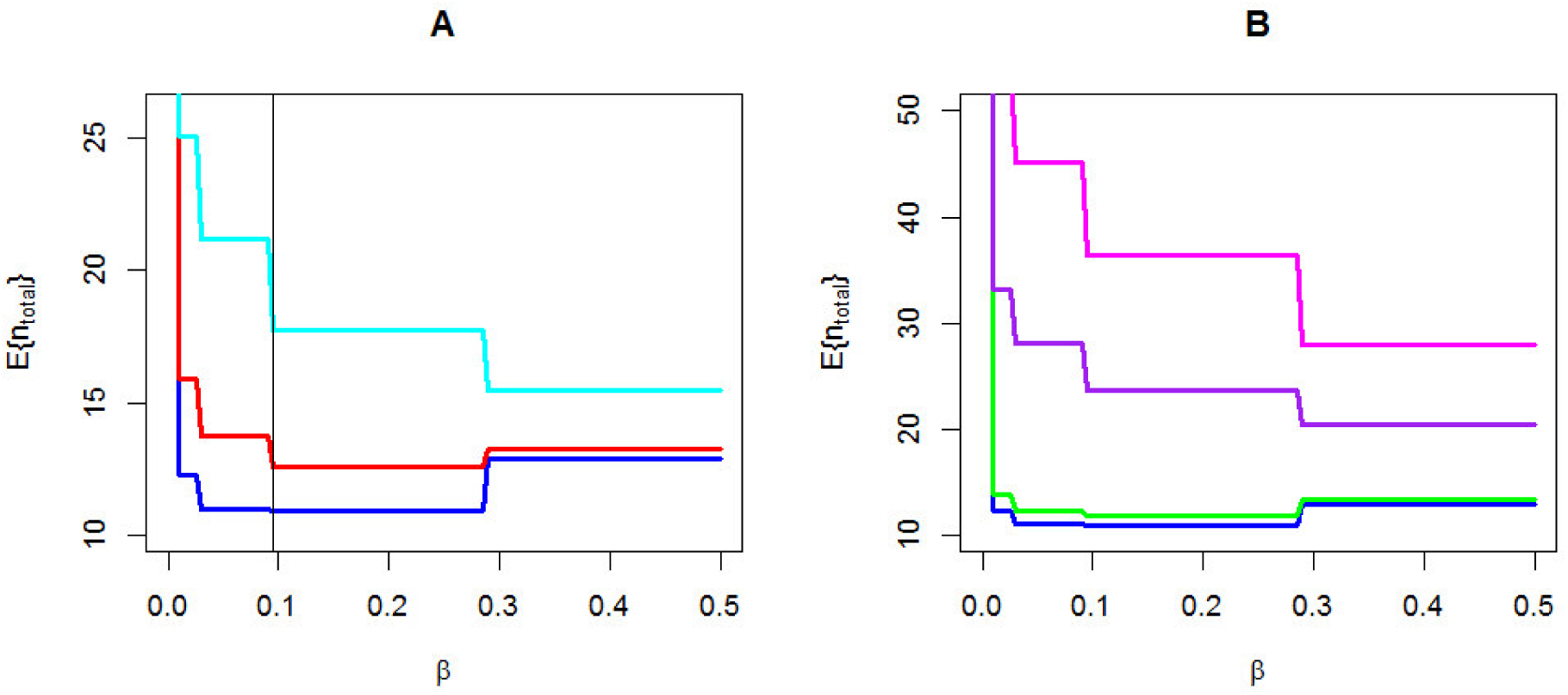
Total number of samples as function of β (= one-power) While E{n_total_} could be depicted as a continuous function of **β**, we present E{n_total_} as a step function. This is due to samples being often indivisible units, such as biological replicates or animals, so that the calculated sample size has to be rounded up to the next integer. Panel A compares different combinations of values for P_pub_ and bias u, whereas Panel B illustrates the consequences of changing the order in which the factors under investigation are examined. In both panels, the ideal scenario is represented in dark blue (P_pub_= 1, u = 0, “correct order”). E{n_total_} is minimized at β_min_ = 0.095 (marked by a black vertical line). Panel A: the red line represents deviation from ideal scenario (P_pub_= 0.5, u = 0.2: only 50% of “negative”/null results are published and 20% excess false positive results were produced by bias). The cyan line represents even more unfavorable conditions (P_pub_= 0.1, u = 0.2) without useful minimum. Panel B: the green line represents a minor error in the testing order, that is, testing the second most probable factor before the first most probable factor. The dark purple line shows the average random order (i.e., all possible testing orders were examined); and the magenta line represents the “worst case,” that is, the inverse testing order. Fixed parameters: π_k_ = 0.5, n_F_ = 10, n_t_ = 10, α = 0.05, δ = 2; n_t_ and n_F_ define the number of teams and the number of investigated factors in the research process, respectively. The sample size was calculated for one-sample one-sided t-tests.

In panel A, the non-monotonic curve progression is very pronounced in the ideal scenario. Therefore, a minimum exists (depicted by the vertical line). Remarkably, its location on the x-axis is independent from the actual parameter combination, although the absolute number of samples at β_min_ differs. As such, any deviation from this ideal scenario leads to an increase of E{n_total_} and, eventually, also to the disappearance of the (useful) minimum.

If pre-study knowledge is not sufficiently exploited, the useful minimum also disappears, which is depicted in panel B. One instance represents a minor error in the experimental order, with small differences from the ideal scenario, whereas another two instances are shown without an existing minimum at all, that is an averaged random and the worst-case inverse order, respectively. Note the different y-axis scale in the two panels. More details are given in the figure caption.

Obviously, any sample size lower than that which minimizes E{n_total_(β)}—being a function of n_a_(β_min_)— must be rejected, since choosing such a sample size would worsen the outcome of the research process in all three respects: E{fp}, E{g}, and E{n_total_}. In other words, a lower power than (1 - β_min_)—in Figure 4, to the right of β_min_—should be avoided.

Higher sample sizes could still be chosen, since this would decrease E{fp} and increase E{g}. However, increasingly higher sample sizes result from only ever smaller gains. Therefore, it is advisable to set a sensible upper limit for n_a_. For example, the maximum efficiency—defined by E{g}/E{n_total_}—can be used to find such limit. Global or local minima of E{n_total_(β)} can then be defined as useful if they are close to the global efficiency maximum.

In conclusion, such useful values for β that minimize the total number of samples exist, when the entire research process is considered. Small deviations from the ideal scenario do not change β_min_. However, these useful minima can be lost, if more unfavorable conditions are assumed. This may, for example, be the case if only a small percentage of “negative”/null results are published or bias leads to some excess of false positive results. Insufficient exploitation of pre-study information can also result in a loss of useful minima.

With regard to these minima (β_min_), our model demonstrates further important facts. The value of β_min_ strongly depends on the quality of pre-study information, π_k,_ and the number of considered factors, n_F_. With higher values of π_k_ and n_F_, the value of (1 - β_min_) also increases. In other words, the use of a higher statistical power in an experiment is especially beneficial if high quality pre-study information is available (and used) and/or more factors considered. The proofs of these propositions are given in the supplement (SI1).

The propositions render it plausible there is a great area in the parameter space of u, P_pub,_ δ, and n_p_ (the number of teams working in parallel) for which useful minima (β_min_) exist. To investigate this question more closely, we varied these parameters and simulated the outcome, which confirmed the assumption. Details are given in the supplementary (SI2). In general, the area of existing useful minima β_min_ in the parameter space reduces if the research community deviates from GSP, but there is still a large such area if the deviations are small.

#### Systematic analysis of the influence of the probability to publish “negative”/null results

The probability to publish “negative”/null results, P_pub_, influences the number of false positive results E{fp} in a complex manner. E{fp} depends on a) the number of non-causal factors that have to be tested before the research process stops and b) the probability for a false positive result per tested non-causal factor. P_pub_ affects both quantities differently.

Considering the probability for a false positive result, false positive results can only be obtained for non-causal factors. If negative test results are published—which is more likely if P_pub_ is high—the number of experiments decreases because other research teams do not have to examine the same factor again. For non-causal factors, this decrease in the number of experiments also decreases the probability to obtain a false positive result (Figure 5, right-hand side).

**Figure 5:**
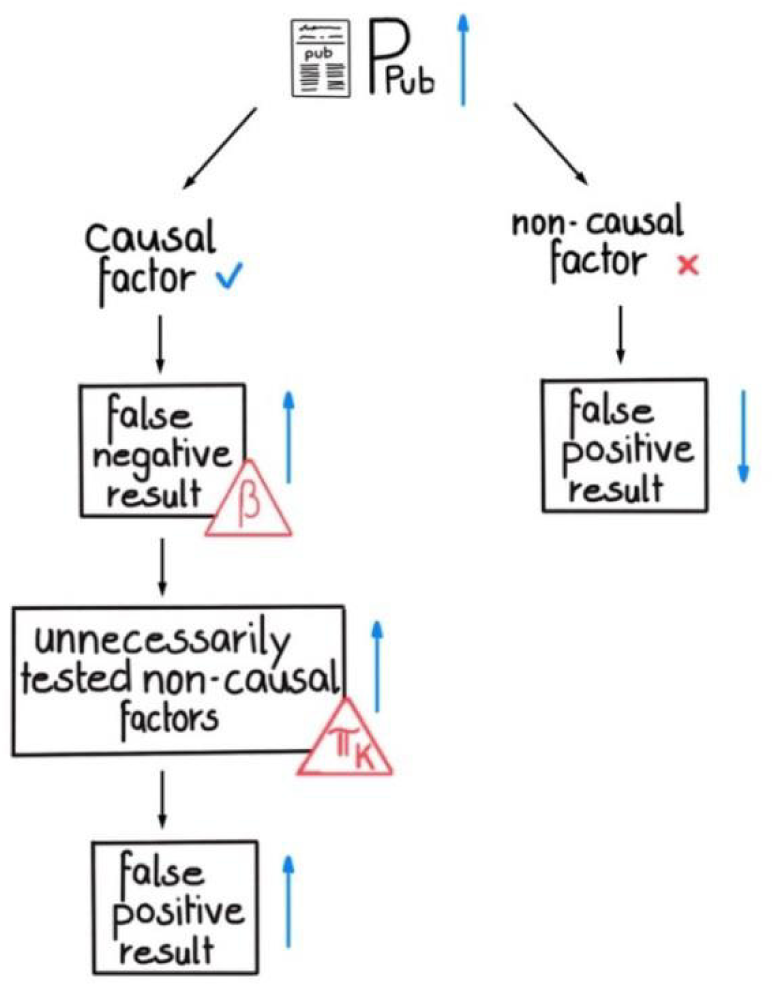
The complex influence of P_pub_ on the number of false positive results. For non-causal factors, an increase of P_pub_ leads directly to a smaller number of false positive results, because the factor is tested less often. For causal factors, an increase of P_pub_ results in a higher number of false “negative”/null results, which strongly depends on statistical power. However, false “negative”/null results lead to unnecessarily tested non-causal factors, and the higher π_k_ is, the higher the probability that the causal factor is tested in the beginning of the research process. A false negative result would therefore have a severe impact on the number of unnecessarily tested non-causal factors. This, in turn, increases the number of false positive results.

For causal factors (i.e., true hypotheses), “negative”/null results can be obtained (depending on β). A higher value of P_pub_ increases the probability that those false “negative”/null results will be published. This, in turn, leads to unnecessary testing of non-causal factors before the research process stops, which again raises the rate of false positive results. This outcome aggravates when the causal factor is tested early in the research process, since more non-causal factors are left to be tested. Hence, the probability for this chance depends on π_k_ (Figure 5, left).

Taken together, this means P_pub_ has contrasting effects with respect to E{fp} on causal and non-causal factors. While a higher P_pub_ leads to a lower false positive rate, it also leads to a larger number of tested non-causal factors. Therefore, the overall effect of P_pub_ on E{fp} depends on which of these contrasting effects is dominant.

Whereas Figure 6 shows E{fp} as functions of β and π_k_ for 11 different publication probabilities, ranging from 0 to 1 (i.e., 0% to 100%), we can show that the publication of all results (P_pub_ = 100%) yields optimal results for most combinations of β and π_k_. Specifically, the topmost area (related to a P_pub_) always represents the lowest expected number of false positives. In the figure, the magenta surface depicts the calculations for P_pub_ = 100%. Although there are also combinations of β and π_k_, for which it seems to be advantageous to not publish negative results (P_pub_ = 0%, black area in Figure 6 A), it is noteworthy that these combinations represent heavily underpowered studies (i.e., studies with insufficient power for given pre-study information and knowledge) and, therefore, are not helpful to the scientific gain. However, there are clear indications that those studies are still published (Button et al. 2013).

**Figure 6:**
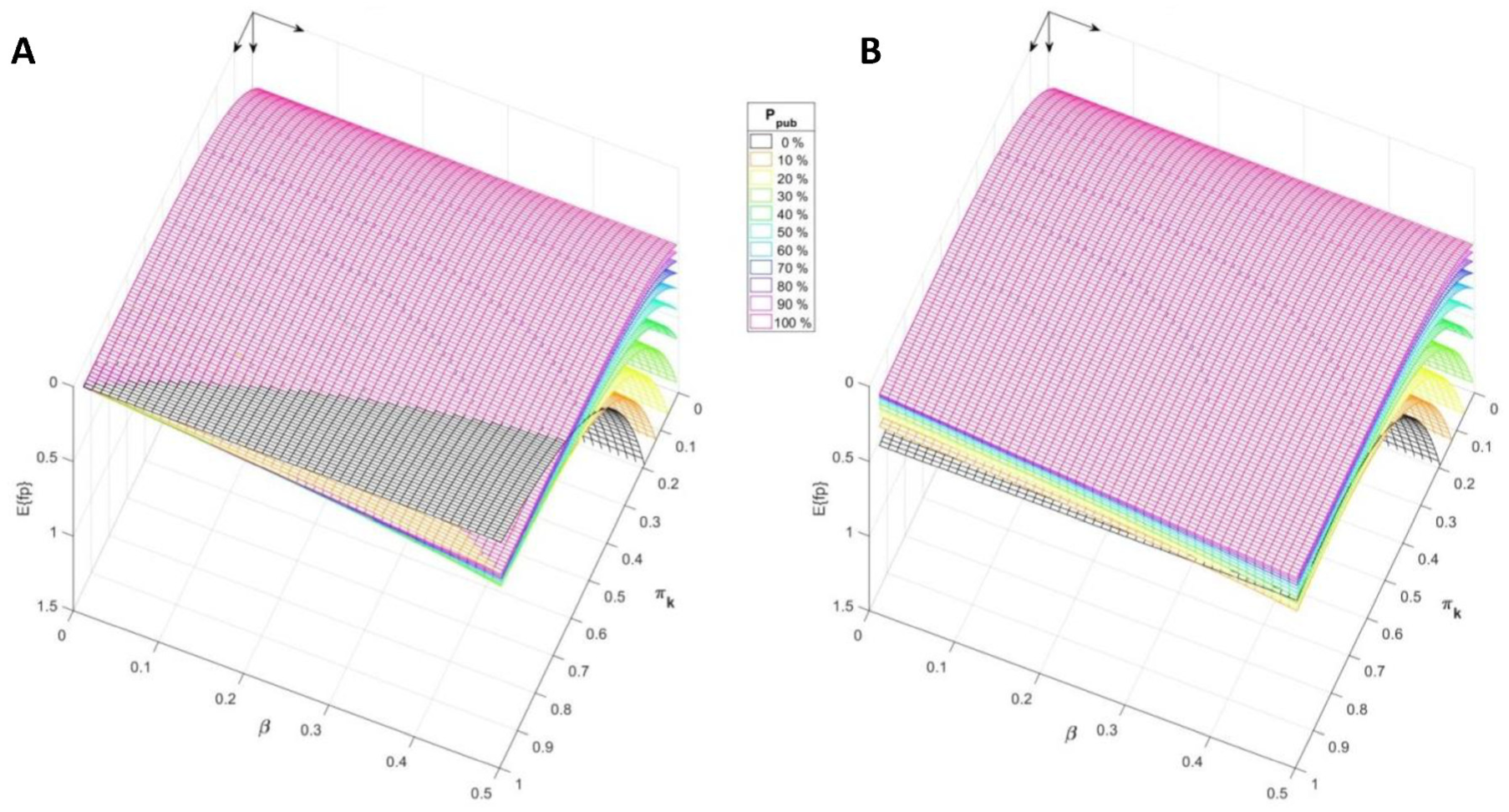
Expected number of false positive results E{fp} as a function of pre-study information, 7t_k_ and P, for 11 publication probabilities. Fixed parameters: n_F_ = 10, n_t_ = 10, a = 0.05, 8 = 2. For a better presentation, the coordinate origin is in the upper rear corner. **Panel A**: The uppermost magenta grid points indicate that, for the largest set of parameter combinations (area ∼ 78%), the publication of all “negative”/null results leads to a favorable lower number of false positives (E{fp}). However, for parameter pairs with higher values of 7i_k_ and p, the uppermost black grid points (P_pub_ = 0%) indicate that any publication of “negative”/null results can be adverse regarding the number of false positives. However, studies represented by this area are not helpful (see text). For publication probabilities between 0% and 100% (indicated by other colors), the statement remains the same. However, the area of ostensible beneficial effect of omitted publications is even smaller. **Panel B** shows the situation when minor errors in the testing order exist, that is, the exchange of the first and second most probable factors. Additionally, this leads to significant increases of the expected number of false positives. There are no combinations of parameters left for which omitting publication is beneficial.

To examine the influence of utilizing pre-study knowledge, we simulated small errors in the testing order of the hypotheses (Figure 6 B). Consistently, if pre-study information is not entirely exploited, it is always better to publish all results to avoid false positive results (only the magenta area is topmost for all values of β and π_k_). The impact of P_pub_ on the total number of samples is similar to the impact on the expected number of false positive results.

### Discussion

We presented a model of the scientific process to show that published results from scientific studies may have an impact on the direction of research, that is, false “negative”/null results can lead to unnecessary (follow-up) experiments. For modeling, it was assumed that a research problem is solved by testing several alternative hypotheses consecutively. Our initial model, describes the relationship between sample size, power, significance, and the expected number of false positives, and was subsequently extended to include any number of research teams, alternative hypotheses, and different probabilities to publish “negative”/null results. Therefore, the probability and effect of publishing false “negative”/null results were included. Additionally, several adjustments were added to reflect deviations from GSP.

This extended model was used to answer two questions: 1) is it possible to reduce the overall consumption of resources by increasing the sample size in individual experiments and 2) under which circumstances and to what degree is the publication of “negative”/null results beneficial? To the best of our knowledge, the presented model considers all relevant elements and parameters necessary to address these questions. For the first question, the model calculations demonstrate that greater sample sizes can lead to a reduction in the overall resource consumption. For animal experiments, this is equivalent to a reduction in the overall number of laboratory animals. If deviations from GSP are just moderate (see below), the statistical power that minimizes the resource consumption (E{n_total_}) is above 80% (i.e., β < 20%) for most of the tested scenarios (Figure 4 and Figure 6).

With respect to the second question, the model shows that not publishing “negative”/null results has always detrimental impacts on the overall resource consumption and the produced excess of false positive results. Only scenarios characterized by a combination of high pre-study probability and low statistical power, in which the reliability of the results is already limited, are exempted. The conclusion derived from these results is to always publish “negative”/null results. This is in line with research results of Nissen et al. (Nissen et al. 2016), but contradicts those of McElreath and Smaldino (McElreath and Smaldino 2015), who claim “publication bias is not always an obstacle, but may have positive impacts,” based on their model calculations. In contrast to McElreath and Smaldino (McElreath and Smaldino 2015), our model considers that an unpublished study to test a hypothesis could well be repeated by another research team and, thus, lead to an unnecessary waste of resources.

Nevertheless, the question arises if our results hold if scenarios with more than moderate deviations from our basic model assumption and from GSP are considered. Obviously, for extreme detrimental assumptions, such as a bias in favor of positive results close to 100% (fraud), the publication of “negative”/null results will not have any effect, nor is it advantageous to use higher sample sizes. If all hypotheses are tested in parallel, higher sample sizes in individual experiments also do not lead to the minimization of overall resource consumption. However, they do still result in higher PPV and the second conclusion (i.e., the beneficial effect of publishing “negative”/null results) still holds. If Joyner et al. are right and the current focus of whole domains of biomedical research is inappropriate (Joyner, Paneth, and Ioannidis 2016) (i.e., the pre-study probability of the tested hypotheses is very low), hardly any measure can improve the reproducibility and efficiency of science, including higher sample sizes. However, the publication of “negative”/null results could still help avoid false positive results and unnecessary resource consumption.

An extensive sensitivity analysis, incorporating diverse deviations, demonstrated that our two main conclusions remain valid for most scenarios (for a complete list of examined scenarios, see SI2). The scenario analyses reveal the order of experiments reflecting decreasing pre-study probabilities is of utmost importance for resource consumption. This underscores the relevance of using pre-study knowledge during scientific planning and questions the use of other criteria for research prioritization. We are aware that our model does not reflect all possible research strategies encountered in practice. For example, the classical group sequential trial design that allows interim analysis (Lehmacher and Wassmer 1999) or the use of historical data in a Bayesian framework (De Santis 2007) may help reduce the resource consumption, while maintaining statistical error rates.

To examine additional interesting scenarios and adapt models for different research conditions and questions, the model scripts (including documentation) written in R (.R) and Matlab (.m) are provided (SI3).

We identified at least two applications for our model. First, it supports a more stringent way to enforce adherence to GSP. Scientific journal editors may enforce this by requesting a self-declaration from authors regarding adequate study design and disclosure of negative, or more generally, of all results independently of outcome. The second is the improvement of incentive schemes for researchers so that the interests of individual researchers are congruent with the improvement of science. Removing the barrier to publishing negative research findings would be an easy way to improve the science and award scientists the incentive of additional publications. Furthermore, our model could be combined with other models (e.g., evolutionary or based on game theory) that describe the actions of individual scientists, given a specific incentive scheme.

Funding agencies, journals, and employers, such as universities and governmental bodies, are especially in demand to implement appropriate measures. These depend on the current state of science in different research areas.

## Supplementary Information: Equations

### Content

In this supplement, the derivations of formulas for the expectations of the scientific gain, the number of false positives, and the total number of samples needed in the entire research process are presented. For comprehensive explanation, the Monte Carlo simulation of the scenario presented in Fig. 2 (main text) that was used to test the validity of the formulas for the expectations is described first. Subsequently, formulas for the above mentioned expectations are presented for the case that only one factor is tested (n_F_ =1). Then we advance to more comprehensive equations for any numbers of factors taken into consideration for a research problem. From these equations, general properties can be derived. Especially, it can be proven that the higher π_k_, the higher the statistical power should be. The corresponding proof is located at the end of this supplement. In addition, formulas for more than one causal factor are presented. For better orientation, please be referred to the list of all relevant variables presented below.

#### List of model variables

|  |  |
| --- | --- |
| $E\{.\}$ | (statistical) expected value of (.) |
| $F_i$ | potential factors to be tested |
| $n_a$ | sample size in a single experiment |
| $n_F$ | number of factors (hypotheses) considered for a research problem |
| $n_{cF}$ | number of (unknown) causal factors (true hypotheses) |
| $n_t$ | number of research teams |
| $n_K$ | number of needed studies to finish research |
| $n_{total}$ | total number of samples |
| $n_p$ | number factors (hypotheses) tested in parallel |
| $\pi_k$ | conditional pre-study probability (of $F_i$) |
| $P_{pub}$ | probability to publish negative results |
| $u$ | bias |
| $\alpha$ | significance level |
| $\beta$ | false negative rate ($1 - \beta = \text{power}$) |
| $\beta_{min}$ | minimal value of $\beta$ regarding $E\{n_{total}\}$ |
| $\delta$ | standardized effect size |
| $\omega_i$ | objective pre-study probability |

#### Description of the Monte Carlo simulation

In our research scenario (main text, Fig. 2) we assume that n_F_ factors are under consideration (F_1_, …, F_i_, …, F_nF_). One or more of these factors are causal for the research problem under investigation. These causal factor(s) are placed on rank(s) between 1 and n_F_ using information of hypothetical exploratory pilot studies. If F_1_ is causal, then F_1_ = 1, and if F_1_ is non-causal, then F_1_ = 0 and so on. The probability of F_1_ = 1 is given by ω_1_. In the Monte Carlo (MC) simulation, this ranking is randomised in accordance with ω_i_ in each iteration. The resulting rankings are stored in a matrix, which dimensions are determined by the number of factors under consideration (n_F_: columns) and the number of simulation runs (N: rows). If there is only one causal factor, the sum of the matrix elements of column i divided by N gives approximately ω_i_.

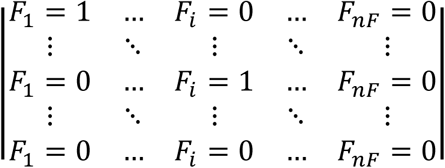

The simulation procedure starts with the first team (team 1) testing F_1_:

If F_1_ = 1, F_1_ is tested positive with the probability 1 — β. In this case and if the number of causal factors n_CF_ = 1, the number of true positives is increased by 1 and the research process stops. If F_1_ is tested negatively (false) this result will be published with the probability P_pub_. If published, all other teams know this negative result and F_2_ is tested next. If the result is not published, team 2 will test F_1_. The testing of F_1_ is repeated until A) one team tests F_1_ positively (we assume that all positive results are published), or B) one team publishes a negative result, or C) all teams have tested F_1_ negatively. Every test conducted increases the number of total samples by the sample size needed for this single experiment.

If F_1_ = 0, the factor is tested positively with the probability α. The number of false positives is then increased by 1. After falsification of the result (an intrinsic characteristic of our simplified scenario) all teams know that the positive result is false and F_2_ is tested next. If F_1_ is tested negatively, it will be published with the probability P_pub_. If published, all other teams know the negative result and F_2_ is tested next. If the result is not published, team 2 will test F_1_. Again, testing of F_1_ is repeated until A) one team tests F_1_ false positively (we assume that all positive results are published), or B) one team publishes a true negative result, or C) all teams have tested F_1_ negatively. Every test conducted increases the number of total samples by the sample size needed for this single experiment.

The process continues until either A) all factors are tested and all teams know the results (either by reading the publication or by testing themselves) or B) all truly causal factors (n_CF_) were tested positively. During the simulation, all true and false positive results and all samples needed in the process are counted. We consider independence among the results obtained from different teams, i.e. false positive and false negative study outcomes are not correlated given the true status (causal/ not causal) of the factor.

It is, however, also possible to derive formulas for the three outputs of the Monte Carlo Simulation, i.e. expectations of the scientific gain, number of false positives, and total number samples. For this, we need to first introduce equations for the pre-study probabilities. These equations are governed by the parameter n_k_, which represents the pre-study information.

Additionally, we introduced a different scenario. Here, more than one factor is tested in parallel by different research teams. The number of factors tested in parallel is denoted by n_p_ and usually np < nF. If all n_p_ test results are published and the truly causal factor is not detected the next np factors will be tested and so on until factors are tested of the truly causal factor is detected. If in contrast a negative test result is not published, one of the teams having all not tested the corresponding factor will repeat this test. As in scenario described above this testing is iterated until one test result is published or all team have tested the factor.

A further extension of the model allows for errors in the validation process. An error of the first kind in the validation means that a false positive result is canonizised, i.e. the process stops with a false result. An error of the second kind in the validation means that the true factor cannot be found.

#### Derivation of equations

##### • Pre-study probabilities

We picture that n_t_ research teams consider n_F_ factors as potential solutions to a research problem, i.e. as causal for an effect under investigation. For simplification, we initially assume that there is only one causal factor (n_cF_ = 1) that explains the observed effect. An exploratory study is performed once which, together with an evaluation procedure, ranks the nF factors according to their likelihood to be causal. Since the outcomes of the exploratory studies can be regarded as random variables, there is an objective pre-study probability that the causal factor is placed on rank i (ω_i_) in this simulation study.

For interpretation of the model outcomes, three properties of ω_i_ are advantageous:

(i) the probabilities should be decreasing (ω_1_ ≥ ω_2_ ≥ … ≥ ω_nF_) i.e. the ranking is optimal,

(ii) the conditional probability (π_k_) for F_i_ = 1 given F_1_ = 0, …, F_i−1_ = 0 should be constant for all i, and

(iii) the asymptotic sum over all ω_i_ should be 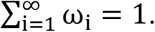

It can easily be derived that:

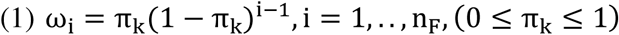

fulfils (i) to (iii). Equation (1) fulfils also (ii) as the conditional probability is given by 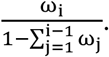

##### • Expectation the total number of samples (E (n_total_); n_F_ = 1)

The only factor considered in this case (F_1_) can be either causal or not. The expected number of studies performed on this factor is the weighted sum of the expected numbers for both cases. Let us first assume that F_1_ is causal. The factor will be tested exactly i times, if the following conditions are fulfilled:

- F_1_ was tested negatively i — 1 times
- none of the corresponding studies were published
- F_1_ is tested positively with the i-th trial or tested negatively with that trial and the study is published.

The maximum number of studies is given by the number of research teams in the field (n_t_). If F_1_ is already tested by n_t_ — 1 teams and none of the test was published, then F_1_ will be tested n_t_ times regardless of the outcome of the experiment.

The expectation of the number of studies for F_i_ = 1 is given by the following series:

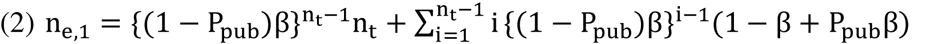

This equation can be simplified using the series 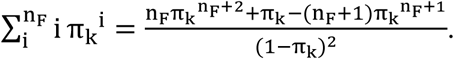

We thus obtain:

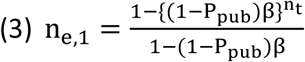

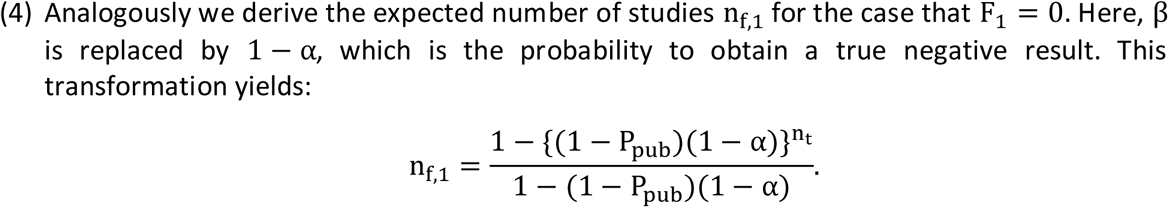

The expected number of studies is then:

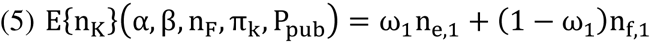

The expectation of the total number of samples used in the research process is given by the product of the number of samples per study n_a_(α, β, 5) and the expectation of the number of studies.

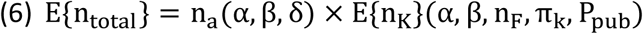

##### • Expectation of the number of false positives (E{fp};n_F_ = 1)

A false positive publication is of course only possible, if F_1_ = 0, which is the case with the probability 1 — ω_1_.

The probability that team i is testing F_1_ positively is given by the probability α multiplied by the probabilities that the preceding i — 1 teams tested F_1_ negatively ((1 — α)^i−1^) and did not publish these results ((1 — P_pub_)^i−1^): α((1 — α)(1 — P_pub_))^i−1^.

The probability that F_1_ is tested positively by any one team is obtained by summing these probabilities over all n_t_ teams (in our model, it is not possible that F_1_ is tested positive by more than one team).

Using the geometric series we thus obtain:

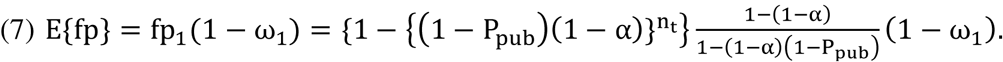

Here 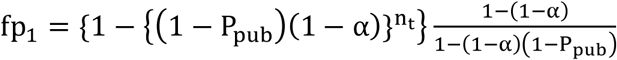 is the expected number of false positives if F_1_ = 0.

##### • Expectation of the number of true positives, i.e. scientific gain (E{g};n_F_ = 1

Similarly we derive the equation for the expected scientific gain, i.e. the number of true positives. Here (1 − α) is replaced by p and 1 − ω_1_ by ω_1_. Therefore

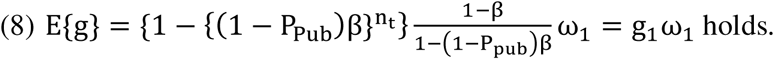

Here, 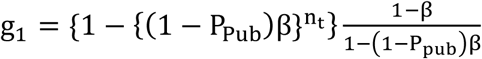 is the expected number of true positives given F_1_ = 1.

### Introduction of bias into the equations

To introduce bias we rely on the approach of Ioannidis (2005) (1) as the “proportion of probed analyses that would not have been “research findings,” but nevertheless end up presented and reported as such, because of bias” (denoted by u)^1^. In equations (3), (4), (7) and (8) β is therefore replaced by β(1 − u) and 1 − α by (1 − α)(1 − u). Thereby we obtain the following equations:

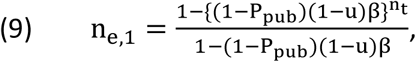

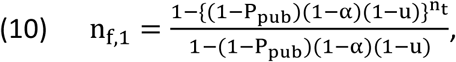

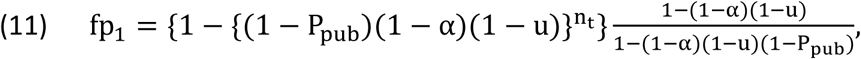

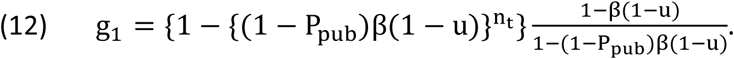

In order to determine the impact of β and α on scientific gain and false positive rate, respectively we differentiate (11) and (12).

The derivation of (11) with respect to α yields:

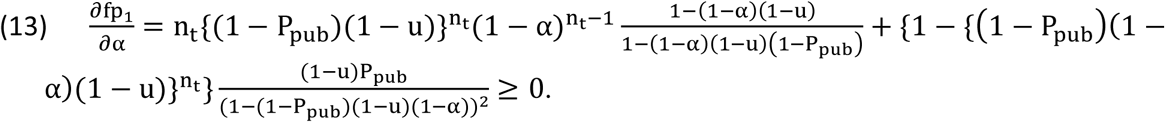

The derivation of (12) with respect to β yields

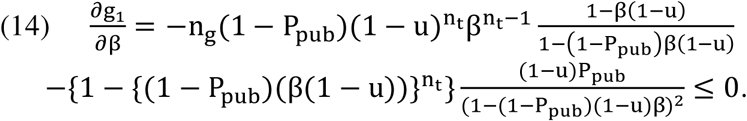

These equations prove that fp_1_ increases with α while g_1_ decreases with β.

### Equations for arbitrary number of factors (n_F_ ≥ 1)

The formula for the total number of samples for any number of considered factors n_F_ is derived by the following consideration:

If F_i_ = 1, which is the case with the probability ω_i_, the mean number of tests of this factor is given by n_e,1_. In this case, the preceding i − 1 non-causal factors were each tested n_f,1_-times in mean. In the case that F_i_ was not recognized as the causal factor, which happens with the probability 1 − g_1_, additional n_F_ − i non-causal factors will be tested n_f,1_-times in mean. In the case that none of the considered factors is causal n_F_ × n_f,1_ studies will be performed in mean. Thus

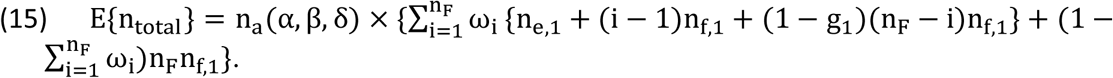

In order to use the parametrization of the pre-study probabilities, we introduce some abbreviations: 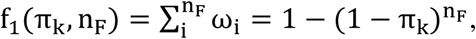 and 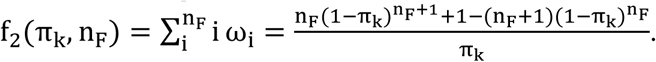

With this notation the expected number of studies in equation (15) can be written as:

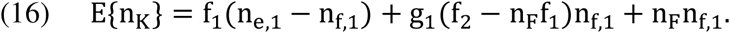

The equation for the expected number of false positives is derived similarly and we obtain:

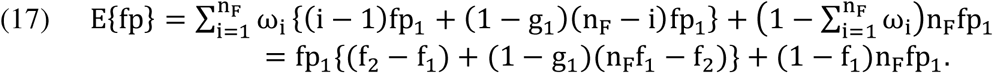

Equation (17) shows that the number of false positives increases with β, because g_1_ decreases with β and g_1_ is the only term in (17) which depends on β. This also means that the absolute number of false positive results decreases with increasing sample size n_a_ and vice versa.

The expectation of the scientific gain is given by the product of the probability that the causal factor is among the n_F_ factors considered 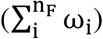 and the probability that a causal factor will be tested positively by one of the n_t_ teams (g_1_):

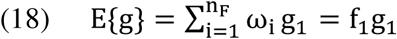

### Parallel testing I

So far, we assumed that the research teams performed the studies consecutively. Let us now consider a scenario in which each factor is tested by all teams in parallel, but in which results are published consecutively. Then n_e,1_ = n_t_ and n_f,1_ = n_t_, while g_1_ and fp_1_ remain unchanged. Therefore, E{fp} and E{g} are unchanged, too, and the following equation is obtained for the expected number of studies:

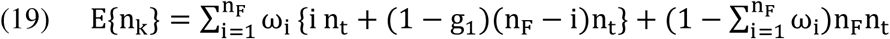

Since in this scenario n_e,1_ and n_f,1_ do no longer depend on P_pub_, publishing negative result would not increase the efficiency of research, i.e. decrease E{n_total_}.

### Parallel testing II (factors are tested in parallel, P_pub_ = 1)

Another possibility is that n_p_ factors are tested in parallel. Then the expectation for the number of studies is:

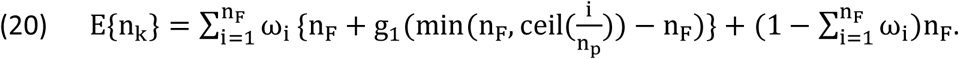

The corresponding expectation for number of false positives is

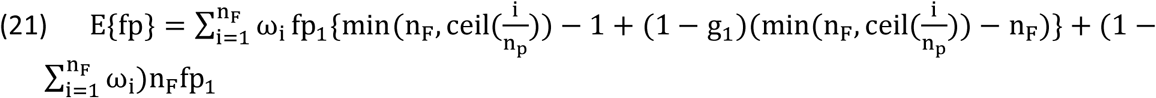

### Equations for arbitrary number of causal factors (n_cF_ ≥ 1)

The model can be extended to the case of more than one causal factor. In this case the research problem is solved, if all n_cF_ causal factors are detected.

In order to derive an equation for E{n_K_} for arbitrary n_cF_, we need an expression for the probability that F_i_ = 1, if there were already n_cF_ − 1 causal factors among the i − 1 factors so far considered. Let us denote this probability by 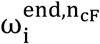.

Furthermore, a formula is needed for the probability P(j; n_F_, n_cF_), that exactly j out of n_cF_ causal factors were among the n_F_ factors.

To obtain a formula for 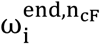, let us consider an ordering of n_F_ factors such that the last of n_cF_ − 1 causal factors is placed on position j, i.e. there are n_cF_ − 2 causal factors on positions preceding j. The probability for such an ordering is 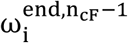. The probability that an additional causal factor is placed on position i > j is given by π_k_(1 − π_k_)^i-(n_cF_−1)−1^. We therefore obtain the following recursion formula:

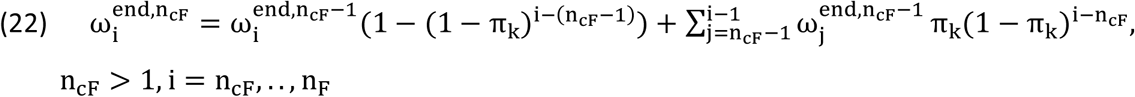

The first summand of (20) takes the case into account that the last of n_cF_ − 1 factors are placed on position i. The term 1 − (1 − π_k_)^i-(n_cF_−1)−1^. describes then the probability that the additional factor is placed on a position from 1 to i − 1. The recursion base of 0 is given by 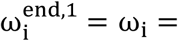 π_k_(1 − π_k_)^i−1^.

To obtain a formula for P(j; n_F_, n_cF_), we consider that there are two ways to choose j out of n_cF_ causal factors among the n_F_ factors.

Either there were j out of n_cF_ − 1 causal factors chosen among n_F_ factors and no additional causal factor will be chosen among the remaining n_F_ − j factors or j − 1 out n_cF_ − 1 causal factors will be selected and one additional causal factor will be chosen among the n_F_ − (j − 1) remaining factors.

Therefore:

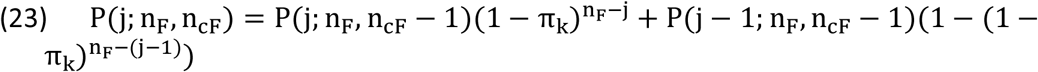

, for j = 1, …, (n_cF_ − l).This recursion formula starts with P(j = 0; n_F_, n_cF_) = ((1 − π_k_)^np^)^ncF^ and P(j = 0; n_F_, n_cF_ = 0) = 1, with P(j; n_F_, n_cF_) = 0 for j > n_cF_ and j < 0.

The equation for E{n_K_} is then derived from the following consideration: If Fj is the last causal factor, exactly n_cF_n_t,1_ + (i − n_cF_)n_f,1_ studies will be performed in mean before the problem is solved, i.e. all n_cF_ factors are discovered. The research problem is not solved with the probability 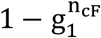. Then additional (n_F_ − i)n_f,1_ studies will be performed in mean. If there were only j < n_cF_ causal factors among the n_F_ considered factors, the expectation of studies performed is equal to (n − j)n_f,1_ + jn_t,1_. Therefore equation (22) follows:

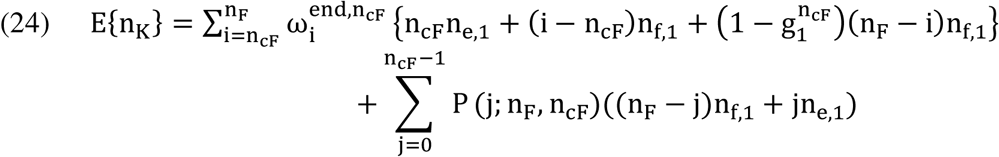

Similarly, it can be shown, that for the number of false positives and the scientific gain

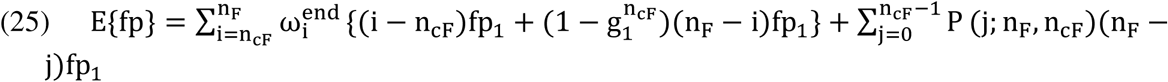

and

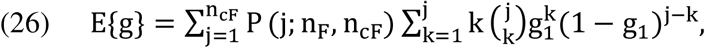

hold, respectively.

### Proof that the probability of a beta error (β_min_) minimizing E(n_total_) decreases with π_k_ and n_F_

We prove that the higher the pre-study probability (π_k_), the higher the recommended statistical power 1 − β and therefore the sample size per study should be. The proof is given here only for n_cF_ = 1. However, as we will see, the proposition depends only on three widely applicable conditions. We further prove that the higher the number of considered factors n_F_, the higher the recommended statistical power 1 − β and therefore the sample size per study should be. This proof could only is only given for P_pub_ = 1, n_cF_ = 1, n_p_=1.

<u>Definition of β_min_:</u>

The smallest probability of an error of the second kind β_m_, which fulfills E{n_total_}(β_m_;π_k_,n_F_,α,δ,P_pub_) ≤ E(n_total_}(β;π_k_,n_F_, α,δ,P_pub_) for all β ∊ [0,1 − α] for fixed π_k_, n_F_, α, δ, P_pub_ is termed β_min_. If n_F_, α, δ, β_pub_ are kept constant, β_min_ is a function of π_k_ denoted by α_min_(π_k_). Similarly the expectation of the number of studies is denoted by E{n_k_}(β,π_k_).

<u>Proposition:</u>

a. From π_k_1__ < π_k_2__ follows β_min_(π_k_2__) ≤ β_min_(π_k_1__).
b. From π_F1_ < ^n^_F2_ follows β_min_(n_F2_) ≤ β_min_(n_F1_).

**Figure S1:** Schematic curves of expectation of total the number of samples per research problem versus ß for two different pre-study probabilities *π*_*k*_1__ and *π*_*k*_2__ (*π*_*k*_1__ < π_*k*_2__). The slope of the curves increase with π_*k*_. Therefore the second curve *E* {n_total_(ß;π_*k*_2__)} is rising at least as long as the first curve *E* {n_total_} (ß; π_*k*_1__) is rising. Then the new minimum can only be found left of the old or at the same position, which is equivalent to the proposition.

*Proof:*

For two different pre-study probabilities π_k_1__ and π_k_2__ there are two different functions of expectation of the total number of samples used for a research problem (illustrated in figure S1). The proposition is proven, if

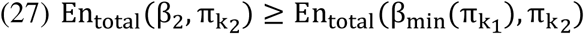

holds for all β_2_ > β_min_(π_k_1__).

We demonstrate that (27) implies the proposition by assuming that β_test_ with β_test_ > β_min_(π_k_1__) is β_min_(π_k_2__). Then inequality (27) requires that En_total_(β_test_, πk_2_) ≥ En_total_(β_min_(π_k_1__), πk_2_), while the definition of β_min_ requires that En_total_(β_test_, π_k_2__) ≤ En_total_(β_min_(π_k_1__), π_k_2__). Both inequalities are satisfied only if En_total_(β_test_,π_k_2__) = En_total_(β_min_(π_k_1__),π_k_2__) .

However, if this is true, then β_test_ is not the smallest β minimizing En_total_, i.e. β_min_(πk_2_).

For illustration see figure S1. The difference in (25) corresponds to the vertical difference of intersection point 4 and intersection point 3 in figure S1.

Fig. S1: Schematic curves of expectation of total the number of samples per research problem versus ß for two different pre-study probabilities *π*_*k*_1__ and *π*_*k*_2__ (π_*k*_1__ < π_*k*_2__). The slope of the curves increase with *π_*k*_. Therefore the second curve *E {n_total_*(ß;*π*_*k*_2__)} is rising at least as long as the first curve *E {n_total_}* (ß; *π*_*k*_1__)* is rising. Then the new minimum can only be found left of the old or at the same position, which is equivalent to the proposition.

To prove (27), we first demonstrate that three relations are sufficient to prove the proposition. We then show that these conditions apply to our model.

The relations are the following:

(i) n_a_(β;α,δ) ≤ n_a_(β_min_; α,δ)(β > β_min_),

(ii) 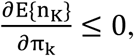 and

(iii) 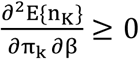

Relation (i) confirms that for higher power a bigger sample size is needed if everything else is kept constant. Additionally, it is assumed that n_a_ is constant throughout the research process. It implies additionally that prior probabilities are not directly used in the sample size determination. Relation (ii) means that we need less studies if our pre-study information is increased. Relation (iii) means that the impact of the statistical power on the number studies is higher, if the causal factors are tested earlier in the research process, which is more frequently the case the greater the pre-study information.

From (i) and the definition of β_min_ follows E(n_K_}(β_2_,π_k_1__) − E(n_K_}(β_min_(π_k_1__)) ≥ 0. From (iii) follows:

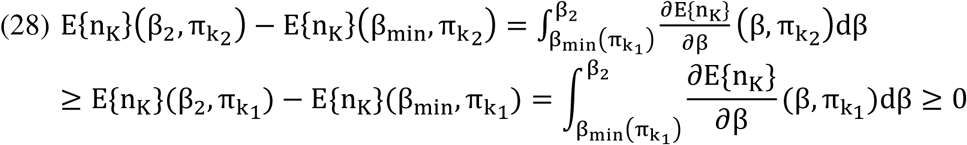

From inequality (26) follows:

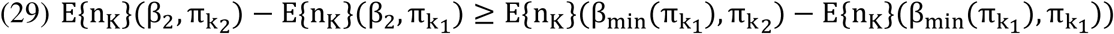

Because of relation (ii) both sides of the inequality (28) are negative. Therefore the following transformation holds:

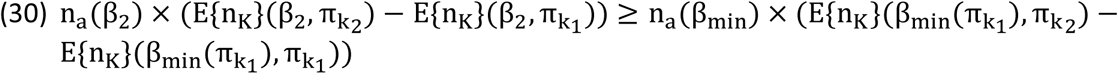

Further transformation yields:

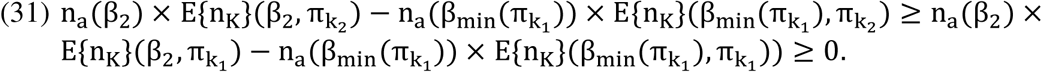

This inequality equivalent to (25) and proves thereby the proposition.

The four intersection points in figure S1 illustrate inequality (29). The left side of the inequality corresponds to the vertical difference of intersection points 4 and 3, the second difference corresponds to that of intersection point 2 and 1. It means that the increase in the total number of samples is even greater from β_min_(π_k_1__) to β_2_, if π_k_ is increased.

We now show that the relations (i), (ii) and (iii) apply to our model:

Relation (i) can be assumed to be the case for all statistical tests. Relation (ii) follows from the differentiation with respect to π_k_ of (16). In equation (16), two terms depend on π_k_:
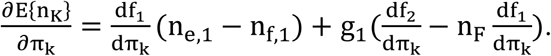

The first term is always negative, because for β ≤ (1 − α) the inequality n_e,1_ ≤ n_f,1_ holds and from the definition of f_1_ follows df_1_/dπ_k_ ≥ 0. For the second term we need to show that 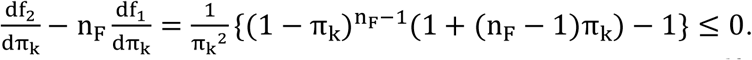

This is done by complete induction. The base clause yields 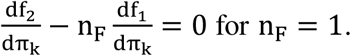
The induction step yields 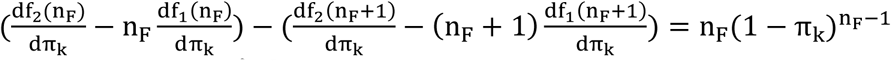
and proves thereby 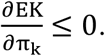

Relation (iii) is equivalent to 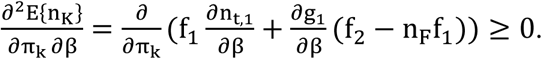

This equation holds, because of 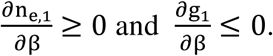

### Properties (ii) and (iii) for parallel testing

It can easily be seen that conditions (i-iii) apply also if parallel testing (I) of one factor by all teams is assumed. The proof can be derived if Eqn. (24) is differentiated with respect to π_k_ and β.

A bit more complicated is the case of n_p_ factors tested in parallel (II). Then the derivation of (25) yields: 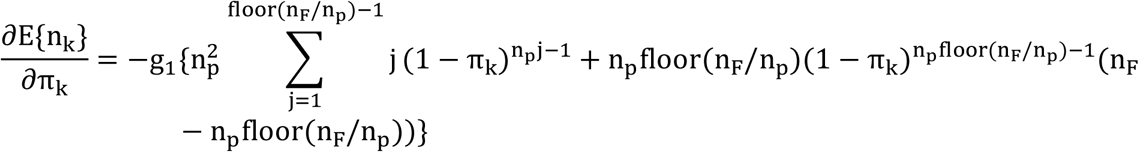

Since all summands in the curly bracket are positive the derivation is negative and condition (ii) holds. Condition (iii) follows then directly from 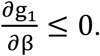

### Proof of proposition b)

We start again with a sufficient condition for proposition b). Such a condition is given by

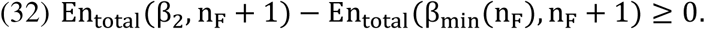

We show that (32) is fulfilled, if

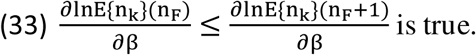

From (33) follows

ln(E{n_k_}(n_F_, β_2_) − ln(E{n_k_}(n_F_, β_min_)) ≤ ln(E(n_k_}(n_F_ + 1, β_2_) − ln(E{n_k_}(n_F_ + 1, β_min_)), which implies:

ln(E{n_k_}(n_F_, β_2_)/E{n_k_}(n_F_, β_min_)) ≤ ln(E(n_k_}(n_F_ + 1, β_2_)/E{n_k_}(n_F_ + 1, β_min_)), and

E{n_k_}(n_F_, β_2_)/E{n_k_}(n_F_, β_min_) ≤ E(n_k_}(n_F_ + 1, β_2_)/E{n_k_}(n_F_ + 1, β_min_).

From the latter inequality and the definition of the minimum 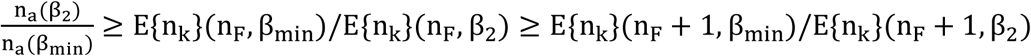 is derived. This inequality implies (32) (given the definition of the minimum).

We now need to demonstrate that (33) applies to the model. Let us assume that P_pub_ = 1. Then 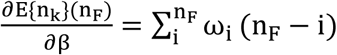
holds.

Since 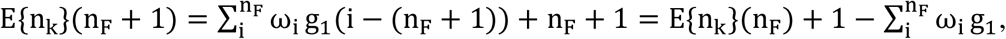, we obtain
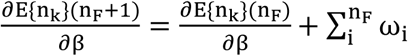 and

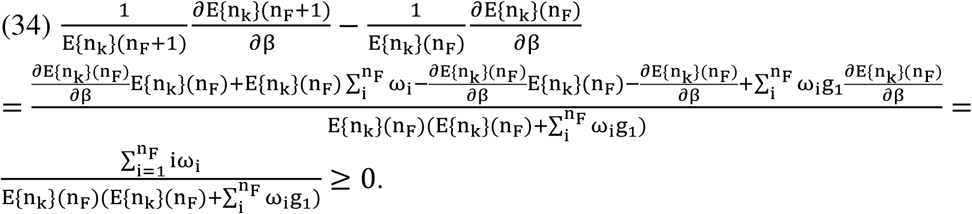

Obviously (34) implies that (33) applies to the model thus proposition b) is valid for the model given P_pub_ = 1.

## Sensitivity analysis

In order to investigate how sensitive our results are when varying the parameters u, P_pub,_ and n_p_ (number of hypotheses tested in parallel),we performed a sensitivity analysis. Additionally, we introduced the possibilities that a positively tested true hypothesis will be falsely rejected in the further research (validation, probability β_v_) or a positively tested false hypothesis will be falsely accepted (probability α_v_).

In this section the following parameters were fixed unless otherwise stated:

δ=2, α=0.05, P_pub_=1, β_v_= α_v_=0, n_p_=1,u =0, n_t_=10 (initial parameter set).

S Fig. 1 shows that for large set of parameter values similar curve shapes result for the total number of samples as function of the probability of a beta-error. S Fig. 2 shows the starting point of the sensitivity analysis: The area where optimization is possible given the initial parameter set. Here, optimizations A) finding a β minimizing the total number of samples and B) maximizing the efficiency E{g}/E{n_total_}. S Fig. 3-8 show that there are less combinations of parameter values (= domain of parameter space) that allow for an optimization as described above if the research community deviates from the good scientific practise. However, a large set of combination remains that allow for an optimization if the deviations are small or moderate.

S Fig. 3 shows the area of the parameter domain in which β_min_ <0.2 for np=1,2,10. From this figure it can be derived that as long as the number of hypotheses tested in parallel (n_p_) is considerably smaller than that of all hypotheses taken into consideration (n_F_). For instance, for n_F_=10 and n_p_=1in all cases where π_k_>0.31 there is β_min_ <0.2, if n_p_=2 β_min_ <0.2 holds for πk>0.39. If n_F_=40 and n_p_=1 leads to β_min_ <0.2 for all π_k_>0.08, for n_p_=2 β_min_ <0.2 for all πk>0.08, and for n_p_=10 β_min_ <0.2 for all π_k_>0.14. Finally, if 100 hypotheses are considered (n_F_=100), then for n_p_=1 and π_k_>0.03 β_min_ is smaller then 0.2, while for np=2 and n_p_=10 the same is true for π_k_>0.04.

S Fig. 4 illustrates the influence of the parameter u, which represents the amount of bias in favor of positive results. It shows that the decreasing size of the area where β_min_<0.2 when u increases u. However, even if u=0.5 the decrease is moderate. For ten considered hypotheses (n_F_=10) in order to obtain β_min_ <0.2 the value of π_k_ must be greater than 0.53. For nF=100 πk>0.06 is required for that purpose instead of π_k_>0.03 for u=0. In contrast for u=0.95 and n_F_=100 π_k_>0.44 is required.

S Fig. 5 similarly shows the same for the parameter P_pub_: while the nonpublication of negative studies results in more samples in general, the impact on the area where β_min_ <0.2 is only slightly decreased, as long at least 50 % of the negative results are published Another possible deviation from the good scientific practice is simply to repeat an experiment if there is a “negative” result. This resembles the questionable research practice described by Bakker (1). The impact of such a procedure is shown in S Fig. 6.

S Fig. 7 illustrates the effect of errors of the first kind in the validation leading to the canonization of false positives. The probability of such an error is denoted by α_v_. The figure demonstrates that the impact of small probabilities α_v_ is small.

S Fig. 8 illustrates the effect of errors of the second kind in the validation leading to the acceptance of false negatives. The probability of such an error is denoted by β_v_. The figure demonstrates that the impact of small β_v_ is small.

**S Fig. 1:**
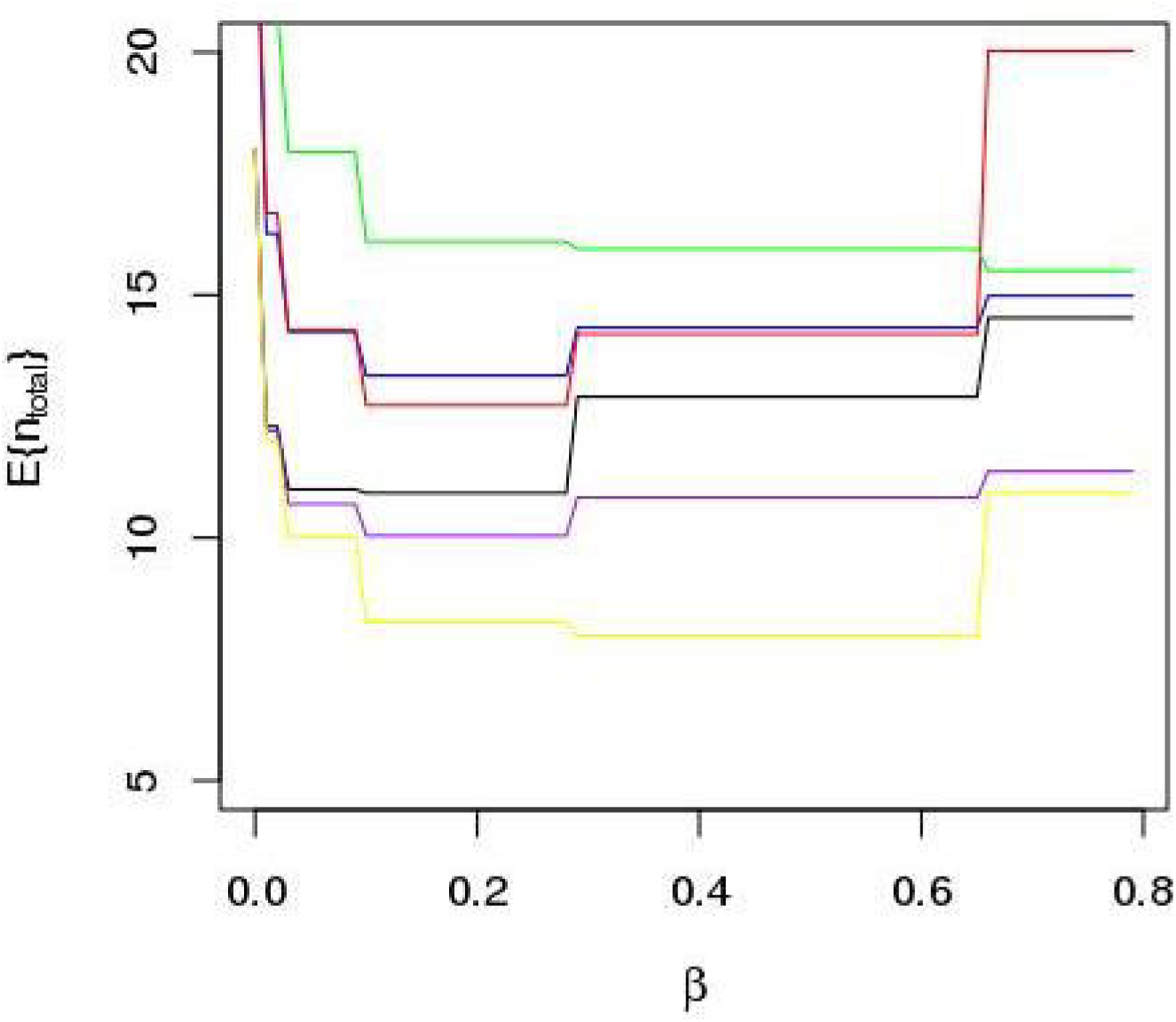
Total number of samples as function of β (=1-power) (nF=10). The black line represents the scenario with no parallel testing, full publication of negative results and no deviation from the good scientific practice. Some additional parameter changes are represented by theother lines:. Scenarios in which more than one scientific hypothesis are tested in parallel by different research teams (n_p_>1) are considered. The scenario where two hypotheses are tested in parallel is represented by the blue line. The scenario where three hypotheses are tested in parallel is represented by the green line. The case of u=0.3 is represented by the purple line. Another scenario of bias (the experiment is once repeated, if the first test result was negative) is displayed by the yellow line. Not publication of negative results in 20% of the cases is represented by the red line.

**S Fig. 2:**
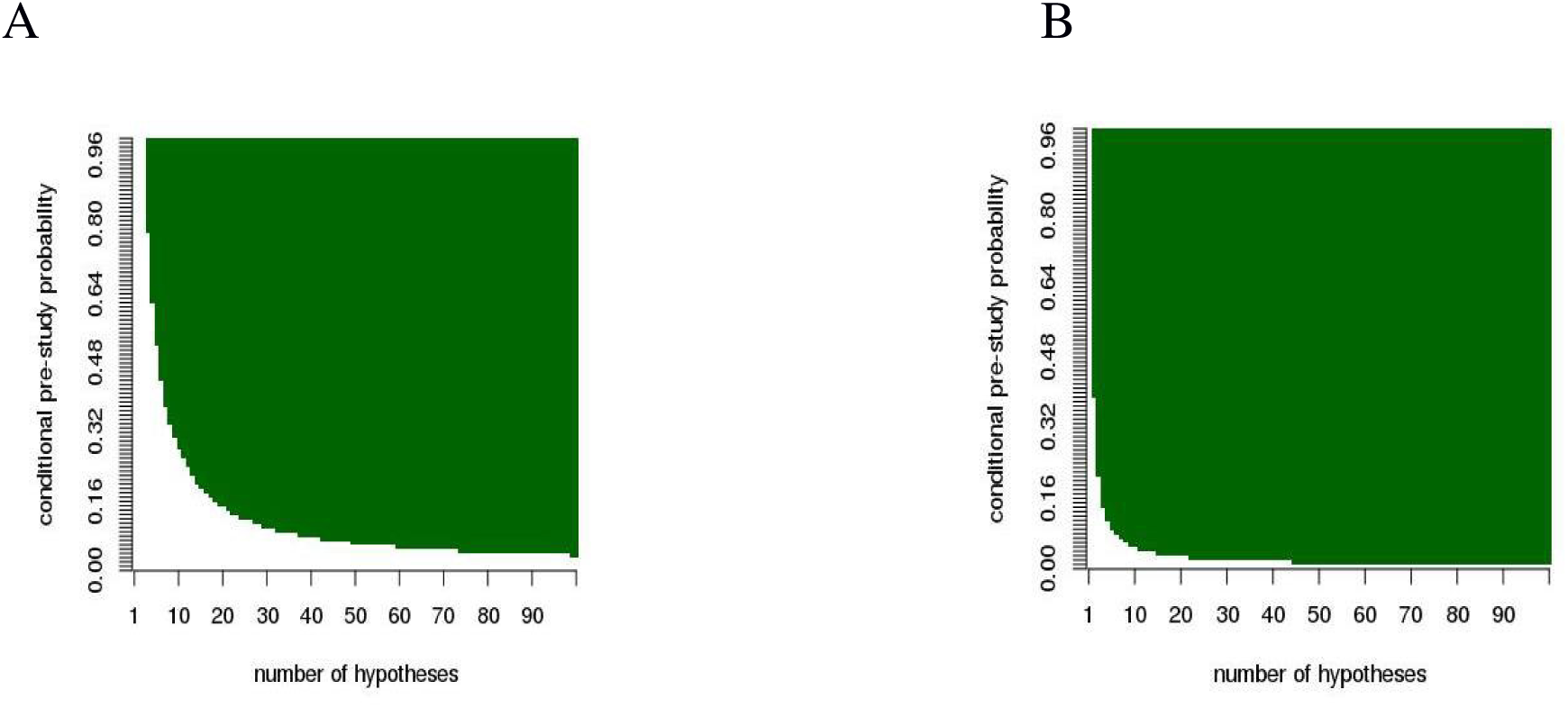
Panel A: Green areas indicate domains of the parameter space in which β_min_ – the probability of an beta-error globally minimizing E{ntotal}- is lower than 0.2, i.e. in this area we find a useful minimum. Panel B : Green areas indicate domains of the parameter space in which β_max_ – the probability of an beta-error globally maximizing E{g}/E{n_total_}- is lower than 0.2.

**S Fig. 3:**
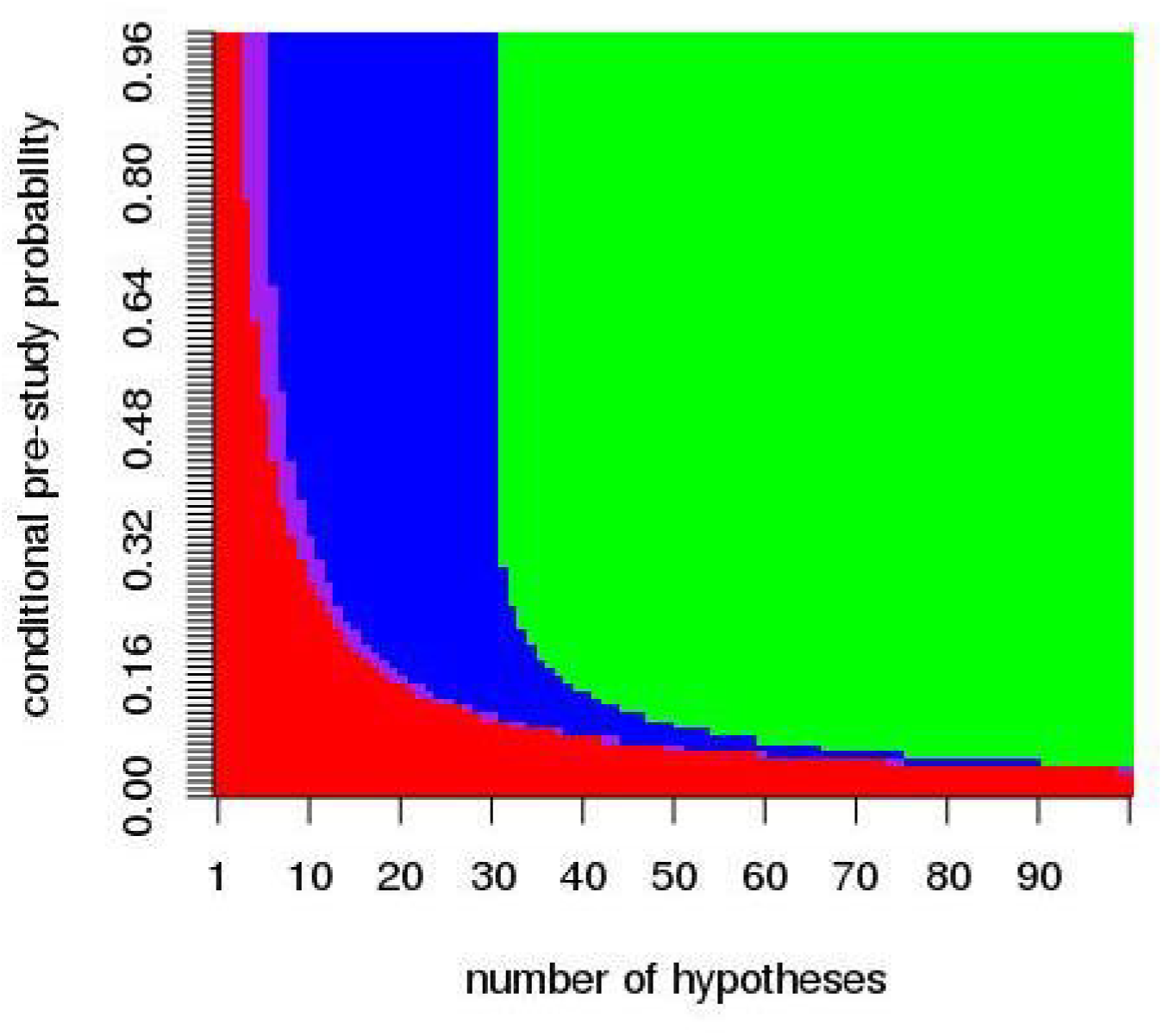
The green area indicates the domain of the parameter space in which β_min_ <0.2, if np =10 (number of hypotheses tested in parallel), the combined area colored green and blue indicate β_min_ <0.2 for np =2, the combination of green, blue, and purple areas (i.e. not the red area) indicate the same for np =1 (basic model assumption).

**S Fig. 4:**
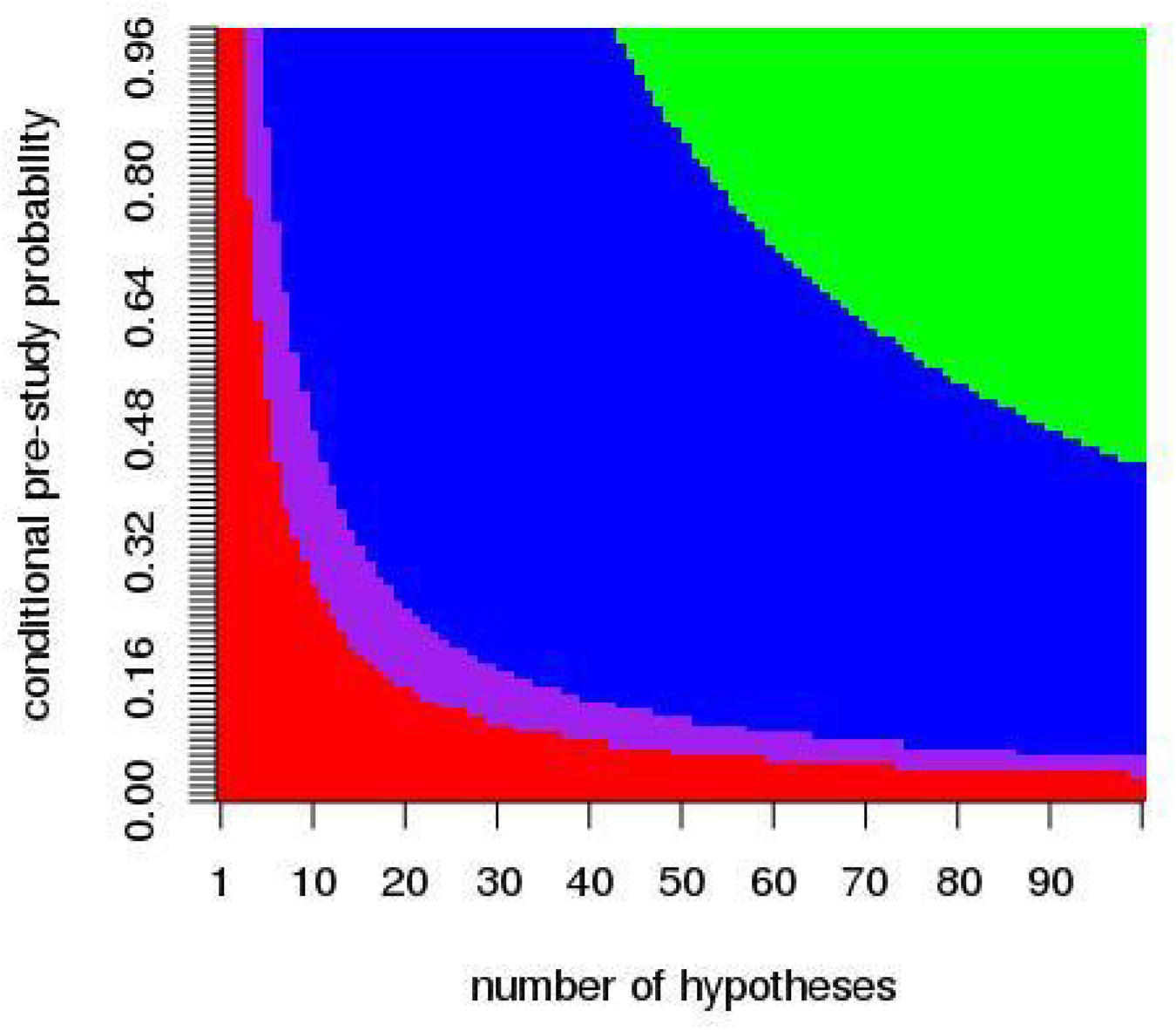
The green area indicates the domain of the parameter space in which β_min_ <0.2, if u=0.95, green and blue areas indicate β_min_ <0.2 for u =0.5, green, blue, and purple for u=0 (basic model assumption).

**S Fig. 5:**
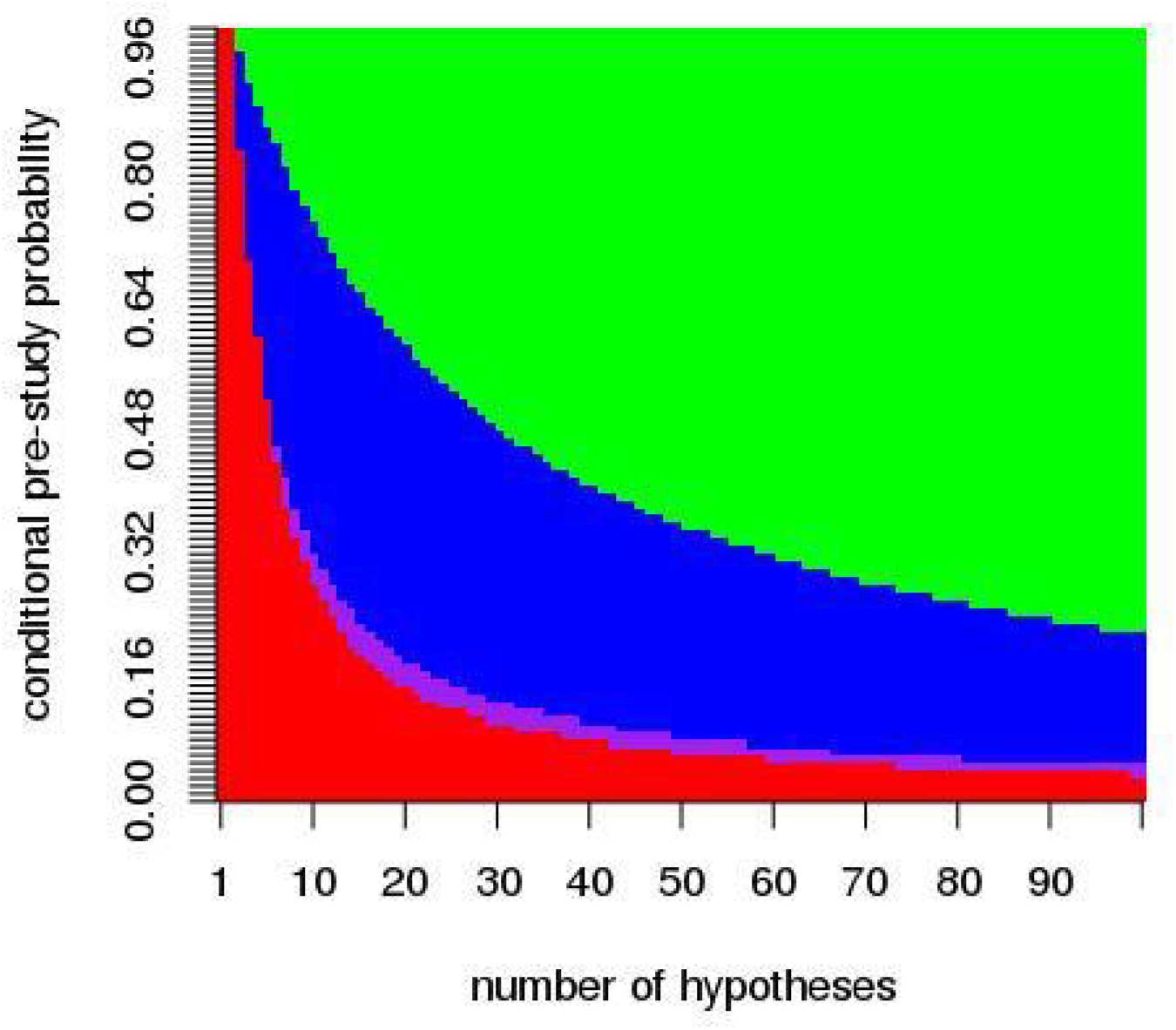
The green area indicates the domain of the parameter space in which β_min_ <0.2, if P_pub_=0.05, the combined area of green and blue indicate β_min_ <0.2 for P_pub_ =0.5, the combination of green, blue, and purple area indicate the same for P_pub_=1, and, finally, the combinations of green, blue, dark blue, and purple areas for P_pub_=1 (basic model assumption). Here are 10 teams assumed (n_t_=10).

**S Fig. 6:**
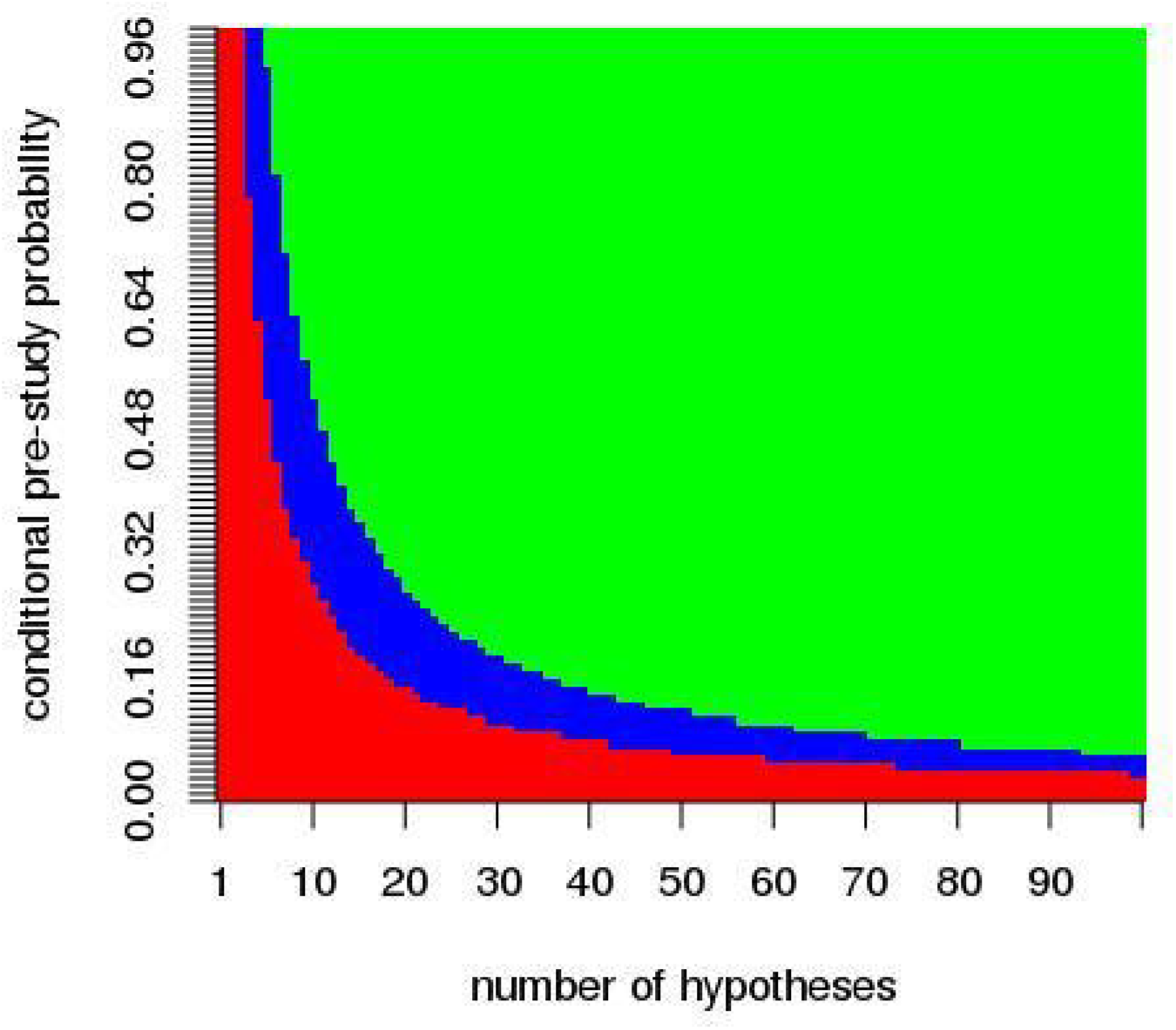
The green area indicates the domain of the parameter space in which β_min_ <0.2, if a negative test is repeated which is an extreme case of the questionable research practice in Bakker et al. (2012) is applied, green and blue areas indicate β_min_ <0.2 if the basic assumptions are applied.

**S Fig. 7:**
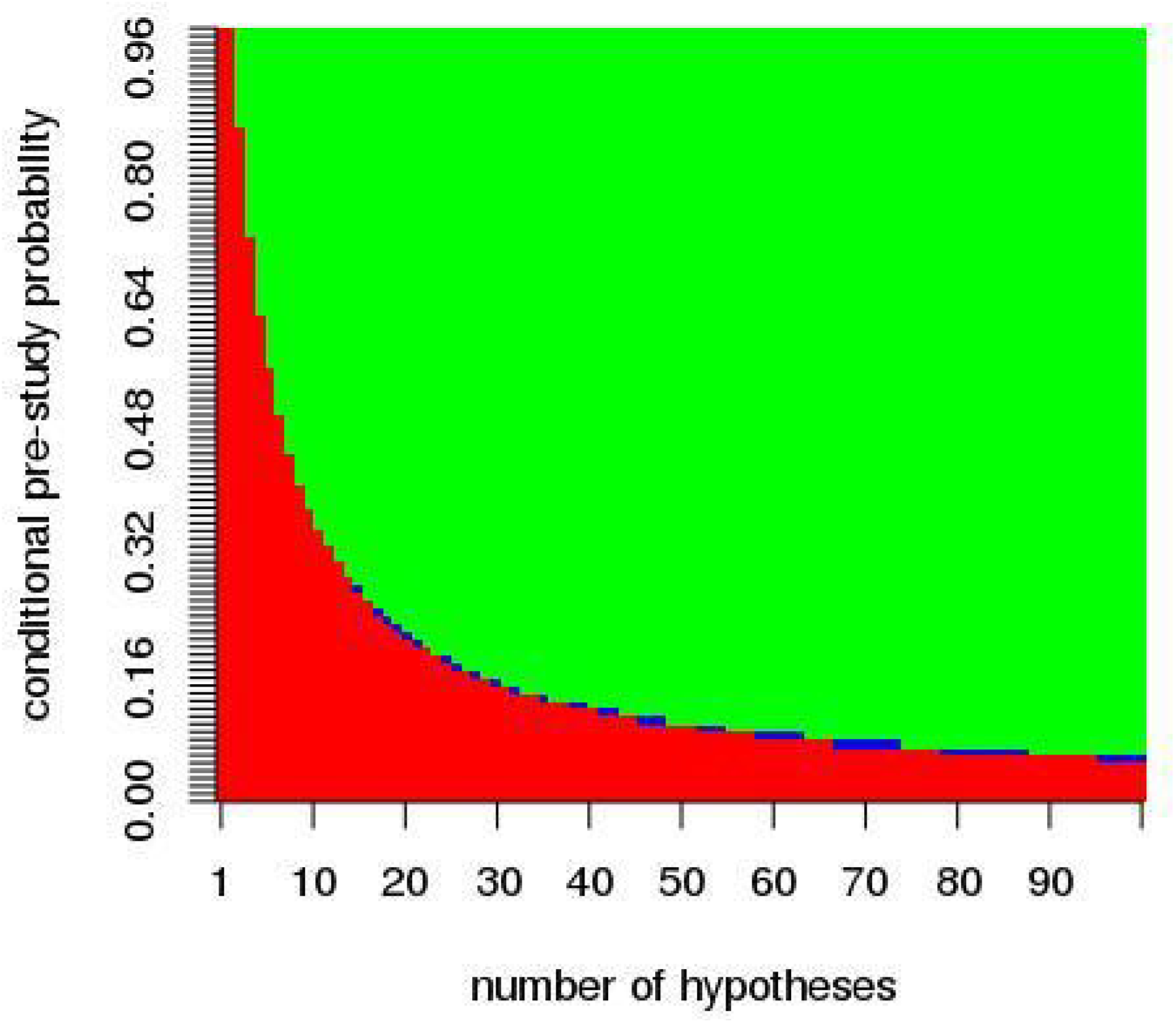
The green area indicates the domain of the parameter space in which β_min_ <0.2, if α_v_=0.05, the combinations of green and blue areas indicate β_min_ <0.2 for α_v_ =0.

**S Fig. 8:**
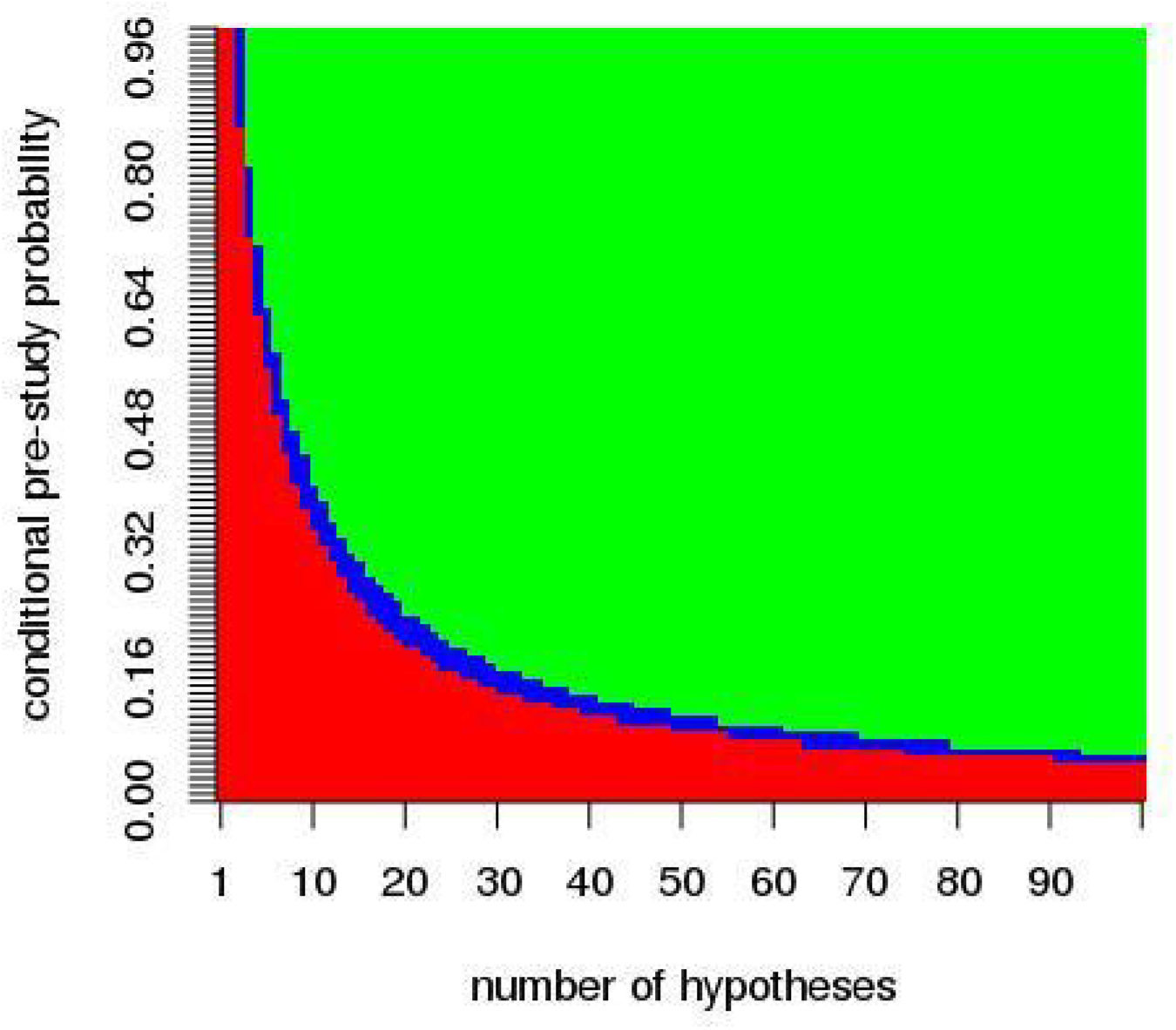
The green area indicates the domain of the parameter space in which β_min_ <0.2, if β_v_=0.05, the combination of green and blue areas indicate β_min_ <0.2 for, if β_v_=0.

## References

Baker, M. 2016. “Is There a Reproducibility Crisis?” Nature 533 (7604):452–454.

Bakker, M., A. van Dijk, and J. M. Wicherts. 2012. “The Rules of the Game Called Psychological Science.” Perspect Psychol Sci 7 (6):543–54. doi:10.1177/1745691612459060.

Bate, Simon T, and Robin A Clark. 2014. The Design and Statistical Analysis of Animal Experiments: Cambridge University Press.

Begley, C. G., and J. P. Ioannidis. 2015. “Reproducibility in science: improving the standard for basic and preclinical research.” Circ Res 116 (1):116–26. doi:10.1161/CIRCRESAHA.114.303819.

Button, K. S., J. P. Ioannidis, C. Mokrysz, B. A. Nosek, J. Flint, E. S. Robinson, and M. R. Munafo. 2013. “Power failure: why small sample size undermines the reliability of neuroscience.” Nat Rev Neurosci 14 (5):365–76. doi:10.1038/nrn3475.

Chambers, C. D. 2013. “Registered reports: a new publishing initiative at Cortex.” Cortex 49 (3):609–10. doi:10.1016/j.cortex.2012.12.016.

Cressey, D. 2015. “UK funders demand strong statistics for animal studies.” Nature 520 (7547):271–272.

De Angelis, C., J. M. Drazen, F. A. Frizelle, C. Haug, J. Hoey, R. Horton, S. Kotzin, C. Laine, A. Marusic, A. J. Overbeke, T. V. Schroeder, H. C. Sox, M. B. Van Der Weyden, and Editors International Committee of Medical Journal. 2004. “Clinical trial registration: a statement from the International Committee of Medical Journal Editors.” N Z Med J 117 (1201):U1054.

De Santis, F. 2007. “Using historical data for Bayesian sample size determination.” J. R. Stat. Soc. Ser. A Stat. Soc 170 (1):95–113.

Erb, Hollis N. 1990. “A statistical approach for calculating the minimum number of animals needed in research.” ILAR J. 32 (1):11–16.

Freedman, L. P., I. M. Cockburn, and T. S. Simcoe. 2015. “The Economics of Reproducibility in Preclinical Research.” PLoS Biol 13 (6):e1002165. doi:10.1371/journal.pbio.1002165.

Head, Megan L., Luke Holman, Rob Lanfear, Andrew T. Kahn, and Michael D. Jennions. 2015. “The Extent and Consequences of P-Hacking in Science.” PLOS Biology 13 (3):e1002106. doi:10.1371/journal.pbio.1002106.

Higginson, A. D., and M. R. Munafo. 2016. “Current Incentives for Scientists Lead to Underpowered Studies with Erroneous Conclusions.” PLoS Biol 14 (11):e2000995. doi:10.1371/journal.pbio.2000995.

Holman, C., S. K. Piper, U. Grittner, A. A. Diamantaras, J. Kimmelman, B. Siegerink, and U. Dirnagl. 2016. “Where Have All the Rodents Gone? The Effects of Attrition in Experimental Research on Cancer and Stroke.” PLoS Biol 14 (1):e1002331. doi:10.1371/journal.pbio.1002331.

Ioannidis, J. P. 2005. “Why most published research findings are false.” PLoS Med 2 (8):e124. doi:10.1371/journal.pmed.0020124.

Ioannidis, J. P. 2006. “Journals should publish all “null” results and should sparingly publish “positive” results.” Cancer Epidemiol Biomarkers Prev 15 (1):186. doi:10.1158/1055-9965.epi-05-0921.

Joyner, M. J., N. Paneth, and J. P. A. Ioannidis. 2016. “What happens when underperforming big ideas in research become entrenched?” JAMA 316 (13):1355–1356.

Kriegeskorte, N., W. K. Simmons, P. S. Bellgowan, and C. I. Baker. 2009. “Circular analysis in systems neuroscience: the dangers of double dipping.” Nat Neurosci 12 (5):535–40. doi:10.1038/nn.2303.

Lehmacher, W., and G. Wassmer. 1999. “Adaptive Sample Size Calculations in Group Sequential Trials.” Biometrics 55 (4):1286–1290.

Marcus, E., and the whole Cell team. 2016. “A STAR is born.” Cell 166 (5):1059–60. doi:10.1016/j.cell.2016.08.021.

McElreath, R., and P. E. Smaldino. 2015. “Replication, Communication, and the Population Dynamics of Scientific Discovery.” PLoS One 10 (8):e0136088. doi:10.1371/journal.pone.0136088.

Munafò, Marcus R., Brian A. Nosek, Dorothy V. M. Bishop, Katherine S. Button, Christopher D. Chambers, Nathalie Percie du Sert, Uri Simonsohn, Eric-Jan Wagenmakers, Jennifer J. Ware, and John P. A. Ioannidis. 2017. “A manifesto for reproducible science.” 1:0021. doi:10.1038/s41562-016-0021.

Nieuwenhuis, S., B. U. Forstmann, and E. J. Wagenmakers. 2011. “Erroneous analyses of interactions in neuroscience: a problem of significance.” Nature Neuroscience 14 (9):1105–1107. doi:10.1038/nn.2886.

Nissen, S. B., T. Magidson, K. Gross, and C. T. Bergstrom. 2016. “Publication bias and the canonization of false facts.” Elife 5. doi:10.7554/eLife.21451.

Simes, R. J. 1986. “Publication bias: the case for an international registry of clinical trials.” J Clin Oncol 4 (10):1529–41. doi:10.1200/jco.1986.4.10.1529.

Smaldino, P. E., and R. McElreath. 2016. “The natural selection of bad science.” R Soc Open Sci 3 (9):160384. doi:10.1098/rsos.160384.

Tsilidis, K. K., O. A. Panagiotou, E. S. Sena, E. Aretouli, E. Evangelou, D. W. Howells, R. Al-Shahi Salman, M. R. Macleod, and J. P. Ioannidis. 2013. “Evaluation of excess significance bias in animal studies of neurological diseases.” PLoS Biol 11 (7):e1001609. doi:10.1371/journal.pbio.1001609.

## References

1. M. Bakker, A. van Dijk, J. M. Wicherts, The Rules of the Game Called Psychological Science. Perspect. Psychol. Sci. 7, 543–554 (2012).

